# Pentose Phosphate Pathway Inhibition activates Macrophages towards phagocytic Lymphoma Cell Clearance

**DOI:** 10.1101/2023.06.09.543574

**Authors:** Anna C. Beielstein, Elena Izquierdo, Stuart Blakemore, Nadine Nickel, Michael Michalik, Samruddhi Chawan, Reinhild Brinker, Hans-Henrik Bartel, Daniela Vorholt, Janica L. Nolte, Rebecca Linke, Carolina Raissa Costa Picossi, Jorge Sáiz, Felix Picard, Alexandra Florin, Jörn Meinel, Reinhard Büttner, Alma Villaseñor, Holger Winkels, Michael Hallek, Marcus Krüger, Coral Barbas, Christian P. Pallasch

## Abstract

Macrophages in the B-cell lymphoma microenvironment represent a functional node in progression and therapeutic response. We assessed metabolic regulation of macrophages in the context of therapeutic antibody-mediated phagocytosis. Pentose phosphate pathway (PPP) inhibition by specific compounds and shRNA targeting induced increased phagocytic lymphoma cell clearance.

Moreover, macrophages provided decreased support for survival of lymphoma cells. PPP inhibition induced metabolic activation, cytoskeletal re-modelling and pro-inflammatory polarization of macrophages. A link between PPP and immune regulation was identified as mechanism of macrophage repolarization. Inhibition of the PPP causes suppression of glycogen synthesis and subsequent modulation of the immune modulatory UDPG-Stat1-Irg1-Itaconate axis. PPP inhibition rewired macrophage maturation and activation *in vivo*. Addition of the PPP inhibitor S3 to antibody therapy achieved significantly prolonged overall survival in an aggressive B-cell lymphoma mouse model.

We hypothesize the PPP as key regulator and targetable modulator of macrophage activity in lymphoma to improve efficacy of immunotherapies.

**Highlights:**

- Macrophage-mediated lymphoma cell phagocytosis is increased by pentose phosphate pathway (PPP) inhibition as an immune regulatory switch for macrophage function and polarization
- PPP inhibition is linked to decreased glycogen synthesis and subsequent modulation of the UDPG-Stat1-Irg1-Itaconate axis
- PPP inhibition is tolerable *in vivo* and facilitates therapeutic targeting of B-cell lymphoma

## Introduction

The tumor microenvironment (TME) represents a hallmark of cancer and interactions between its transformed and non-transformed immune bystander cells determine disease progression and therapeutic response [1]. Tumor cells generate a tumor-supportive milieu by cytokine and metabolite secretion. These mediators alter occurrence of bystander cells and shift the activity of the infiltrating immune cells from an anti to a pro-tumoral response [2]. Particularly tumor associated macrophages (TAMs) exert tumor-supportive functions [3]. TAMs are immunosuppressive towards other tumor infiltrating immune cells and promote tumor growth by enhancing vascularization and metastasis [4][5].

In contrast, we have shown that macrophages, despite their pro-tumoral activity, are central for tumor cell clearance in the context of immunotherapy against aggressive B-cell lymphoma, but their phagocytic capacity becomes impaired by lymphoma cells [6].

The last decades have led to the development of numerous new treatment strategies against B-cell malignancies. These new therapies prolonged patient survival, but have mostly been unable to greatly improve cure rates, especially in chronic lymphocytic leukemia (CLL). The front-line strategy is chemo-immunotherapy, in which antibodies like rituximab, or obinutuzumab are combined with chemotherapeutic agents. Although this combinational therapy achieved improved therapy efficacy, we demonstrated, that leukemia cells in therapy refractory niches are able to diminish the engulfment of antibody-targeted tumor cells by macrophages and thereby decrease therapy efficacy and cause relapse [6].

The function of macrophages is determined by the local environment impaction on their differentiation and polarization. Macrophages are a heterogeneous population with different subtypes exerting pro-inflammatory and phagocytic (M1-like macrophages), and anti-inflammatory and tissue regenerative (M2-like macrophages) activities. TAMs represent a mixture of these two characteristics with a gradient towards the anti-inflammatory and phagocytic inactive phenotype [7][8]. The TME includes activation mediators such as cytokines, chemokines, and metabolites, which control the polarization of contained macrophages. Alterations in the microenvironment initiate changes in macrophage metabolism, which are crucial for polarization and the closely linked macrophage function [9][10]. Changes in the cellular metabolism have the ability to repolarize the macrophage phenotype by which pro-tumoral TAMs could acquire anti-tumoral activity [11][12][13]. The interaction between macrophages and lymphoma cells and the metabolic sensitivity of macrophages open up a new strategy to optimize anti-cancer therapy by modulation of the metabolic activity of macrophages to improve their anti-tumor efficacy and diminish their tumor supportive function.

Here we demonstrate that pentose phosphate pathway (PPP) inhibition in macrophages increases their activity and phagocytic capacity whereby pro-tumoral bystander function is diminished. As driving mechanism, we discovered a new connection between metabolism and immune regulation by modulation of the UDPG-Stat1-Irg1-Itaconate axis. The effects of PPP inhibition were transmitted into human patient samples and also reproduced *in vivo*, where significantly increased survival in an aggressive lymphoma mouse model was achieved. These results open up a promising new treatment strategy against B-cell malignancies in clinical use.

## Results

### 1. Metabolic inhibition of the pentose phosphate pathway leads to increased phagocytic capacity of macrophages

To investigate how metabolic modulation of TAMs in the context of immunotherapy might affect phagocytic capacity, we performed a metabolic focused screening approach for antibody-dependent cellular phagocytosis (ADCP). Key metabolic pathways were blocked using representative inhibitors in a macrophage and humanized aggressive B-cell lymphoma (hMB) co-culture-assay system and phagocytosis was assessed through specific antibody targeting (alemtuzumab; anti-CD52) (Fig.1A). Approaching inhibition of glycolysis (via 2-deoxy-D-glucose), AMP-activated protein kinase (AMPK) mediated cell energy regulation (via BML-275), mitochondrial ATP production (via oligomycin), and the pentose phosphate pathway (PPP) (via 6-aminonicotinamide and oxythiamine), the metabolic ADCP screening was conducted using non-toxic concentrations (Suppl.Fig.1). The metabolic inhibition was performed in both, by treatment of the co-culture and by pre-treatment of each cell type (macrophage or lymphoma cell), to infer specific macrophage vs. lymphoma cell phagocytic interactions.

**Fig.1.**
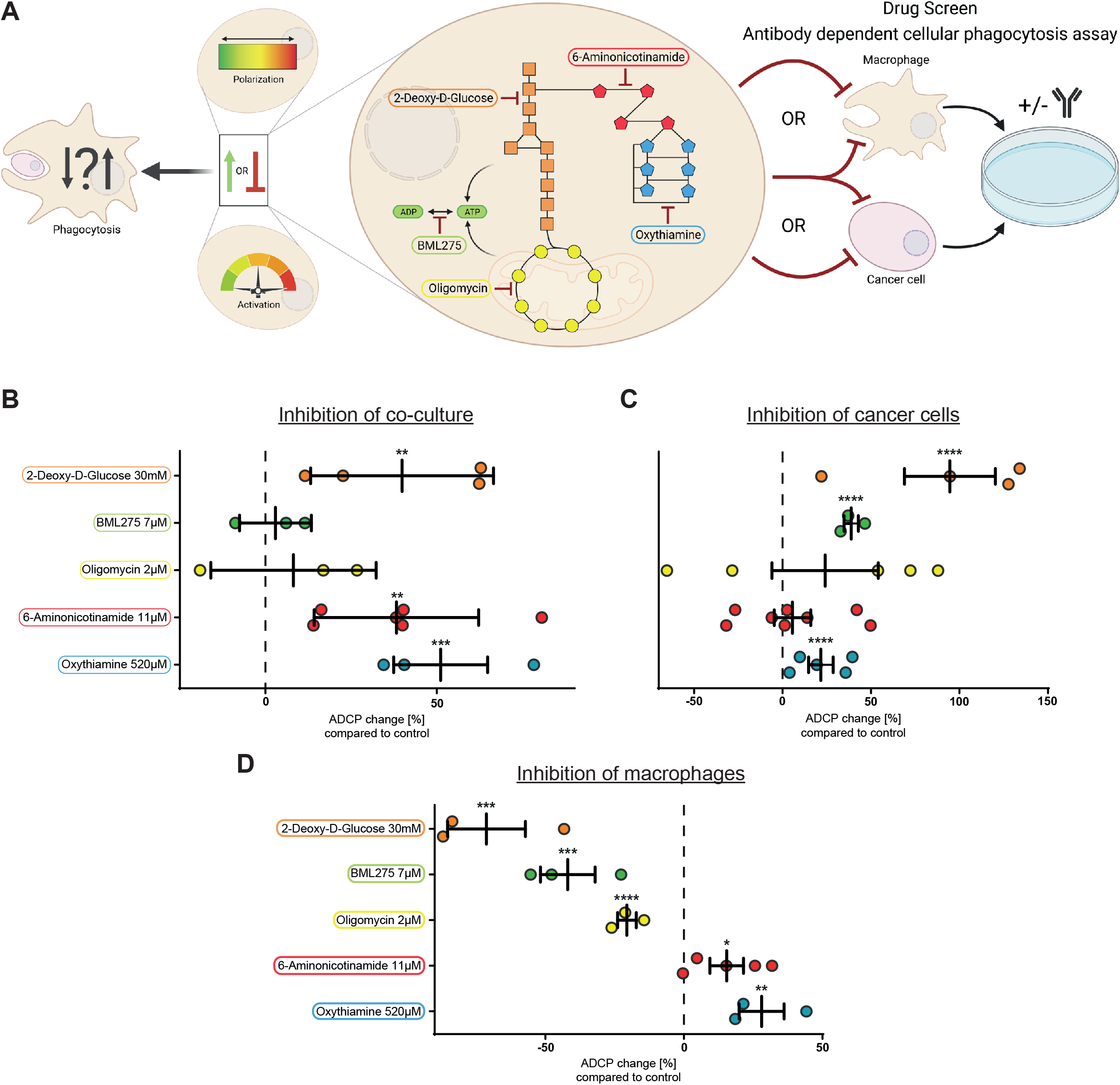
Metabolic inhibition of pentose phosphate pathway leads to increased phagocytic capacity of macrophages. **(A)** Scheme of ADCP-based metabolic screening approach. **(B-D)** ADCP change compared to basal phagocytosis rate of J774A.1 macrophages under inhibition of respective metabolic pathways. **B** Inhibition of all co-culture components, **C** Inhibition of only hMB cells, **D** Inhibition of only macrophages. n=3-6. Data are shown as mean ± SEM. *P* values were calculated using unpaired t-test. \**p* < 0.05; \*\**p* < 0.01; \*\*\**p* < 0.001; \*\*\*\**p* < 0.0001.

Glycolysis inhibition significantly increased ADCP rates in co-culture (+40%, *P*<0.01) and by pre treatment of lymphoma cells (+95%, *P*<0.0001), while pre-treatment of macrophages significantly diminished ADCP rate (-71%, *P*<0.001) (Fig.1B-D). Similarly, AMPK inhibition increased ADCP rate significantly in the lymphoma pre-treatment setting (+39%, *P*<0.0001), while a significantly diminished ADCP rate was seen by pre-treatment of macrophages (-42%, *P*<0.001) (Fig.1C-D). Inhibition of mitochondrial ATP production also diminished ADCP rate significantly in the macrophage pre-treatment setting (-21%, *P*<0.0001) (Fig.1D).

Solely inhibition of the PPP induced significantly increased ADCP rates when treatment was added to the co-culture or macrophage were pre-treated. The increase was induced by both, inhibition of the oxidative part of the PPP via 6-phosphogluconate dehydrogenase inhibition (6Pgd; inhibitor 6 aminonicotinamide) (co-culture +40% *P*<0.01; macrophage pre-treatment +15%, *P*<0.05) as well as inhibition of the non-oxidative part *via* transketolase inhibition (Tkt; inhibitor oxythiamine) (co culture +51% *P*<0.001; macrophage pre-treatment +28%, *P*<0.01) (Fig.1B, D). Moreover, pre treatment of lymphoma cells with oxythiamine increased phagocytic rate significantly (+19%, *P*<0.0001).

Of note, Tkt inhibition induced the highest increase in phagocytic capacity in the co-culture and by pre-treatment of macrophages in the screening approach.

Thus, inhibition of glycolysis, AMPK, and mitochondrial ATP production negatively affected macrophage phagocytic capacity, while blocking PPP favoured tumor clearance by macrophages.

### 2. Cross validation of PPP inhibition in macrophages confirms increased ADCP capacity

To further investigate the PPP in context of macrophage function and as a target for improving immunotherapy we applied alternative inhibitors for the oxidative and the non-oxidative part of the PPP (6Pgd: physcion, Tkt: p-hydroxyphenylpyruvate) [14][15] and confirmed significant increases in ADCP rates (physcion +26%, *P*<0.0001; p-hydroxyphenylpyruvate +25%, *P*<0.001) (Fig.2A). Additionally, we recapitulated the phagocytosis assays with the human monocyte cell line THP1 using an alternative antibody (obinutuzumab; anti-CD20 Type II), also identifying significant induction of ADCP under PPP inhibition (6-aminonicotinamide *P*<0.05; oxythiamine *P*<0.01) (Fig.2B).

**Fig.2.**
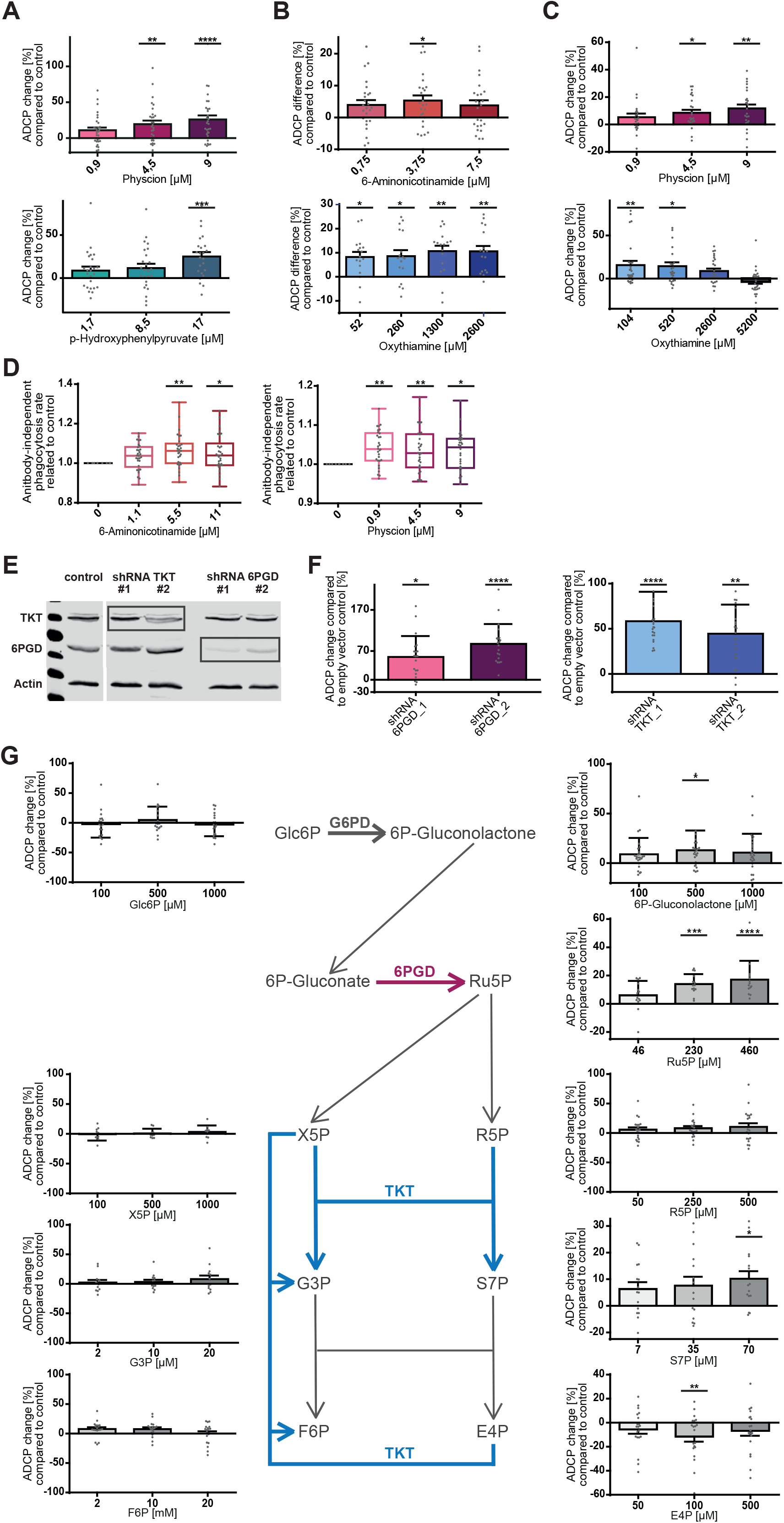
Cross validation of PPP inhibition in macrophages confirms induction of increased ADCP capacity. **(A-C)** ADCP change compared to basal phagocytosis rate of J774A.1 macrophages under inhibition of PPP. **A** Alternative inhibitor physcion of 6-phosphogluconate dehydrogenase (6Pgd) in oxidative part of PPP (red) and p-hydroxyphenylpyruvate for inhibition of transketolase (Tkt) in non-oxidative part of PPP (blue). **B** Using human monocyte cell line THP1 and CD20 antibody obinutuzumab under inhibition of 6Pgd by 6-aminonicotinamide (red) and inhibition of Tkt by oxythiamine (blue). **C** ADCP assay performed under hypoxic conditions and inhibition of 6Pgd by physcion (red) or inhibition of Tkt by oxythiamine (blue). **(D)** Antibody-independent cellular phagocytosis of hMB cells by J774A.1 macrophages compared to control under inhibition of 6Pgd by 6-aminonicotinamide (left) and physcion (right). **(E)** western blot of J774A.1 macrophages transfected with shRNA empty vector control, shRNA targeting Tkt, and shRNA targeting 6Pgd. **(F)** ADCP change compared to basal phagocytosis rate of empty vector control J774A.1 macrophages under shRNA mediated knockdown of 6Pgd (red) and Tkt (blue). **(G)** ADCP change compared to basal phagocytosis rate of J774A.1 macrophages under supplementation of metabolites of the PPP. Enzyme reactions in focus coloured in violet (6Pgd) and blue (Tkt). *E4P* erythrose-4-phosphate *, F6P* fructose-6-phosphate, *G3P* glyceraldehyde-3-phosphate, *Glc6P* glucose-6-phosphate, *R5P* ribose-5-phosphate, *Ru5P* ribulose-5-phosphate, *S7P* sedoheptulose-7-phosphate, *X5P* xylulose-5-phosphate. n=3-6. Data in **A-D, F-G** are shown as mean ± SEM, in **E** one representative western blot example. *P* values were calculated using one-way ANOVA. \**p* < 0.05; \*\**p* < 0.01; \*\*\**p* < 0.001; \*\*\*\**p* < 0.0001.

Since hypoxia is a functional aspect of the TME *in vivo*, we also conducted ADCP assays under hypoxic conditions (O _2_ 1,5%) and observed significantly increased ADCP rates under inhibition of the oxidative and the non-oxidative part of the PPP (physcion +12%, *P*<0.01; oxythiamine +20%, *P*<0.01) (Fig.2C).

To evaluate if PPP inhibition also increases phagocytic capacity of macrophages without the targeting function of antibodies, we assessed antibody-independent cellular phagocytosis (AICP) (Fig.2D) and observed significantly increased AICP rates only by inhibition of the oxidative part of the PPP (6 aminonicotinamide *P*<0.01, physcion *P*<0.01).

Lastly, to further assess the impact of 6Pgd and Tkt on ADCP capacity and to abrogate off-target effects of inhibitors, we generated shRNA knockdown for the respective enzymes in macrophages (Fig.2E, Suppl. Fig.3P-Q). Silencing of 6Pgd and Tkt significantly increased macrophages ADCP (6Pgd +87%, *P*<0.0001; Tkt +58%, *P*<0.0001) (Fig.2F).

Altogether, we demonstrate that PPP enzyme inhibition by metabolic inhibitors as well as 6Pgd and Tkt knockdown in macrophages promotes phagocytosis of lymphoma cells and warrants further investigation of the molecular function.

### 3. Increased phagocytosis is driven by PPP enzyme inhibition and not by PPP metabolite shifting

To identify which specific components and metabolites of the PPP directly affect phagocytic function, we performed phagocytosis assays with supplementation of single educts and products of the PPP (Fig.2G). We observed unaltered ADCP rates by the non-exclusive PPP metabolites glucose-6-phosphate [Glc6P], ribose-5-phosphate [R5P], xylulose-5-phosphate [X5P], glyceraldehyde-3-phosphate [G3P], and fructose-6-phosphate [F6P]. In contrast, supplementation of the PPP exclusive glucose-6-phosphate dehydrogenase (G6pd) product 6-phosphogluconolactone significantly increased ADCP rate (+13%, *P*<0.05) (Fig.2G; right panel). Addition of the PPP exclusive metabolites ribulose-5-phosphate [Ru5P] (product of 6Pgd) and sedoheptulose-7-phosphate [S7P] (product of Tkt) respectively, significantly increased ADCP rates (Ru5P +17%, *P*<0.0001; S7P +20%, *P*<0.05) (Fig.2G; right panel). In contrast, supplementation of the Tkt educt erythrose-4-phosphate [E4P] significantly reduced ADCP rate (-12%, *P*<0.01) (Fig.2G; right panel).

In conclusion, we identified products of Tkt and 6Pgd to promote macrophages’ phagocytic activity while enzyme educts are diminishing macrophage activity. This indicates that the inhibition of the enzymes itself and not a decrease in their products leads to the increased phagocytic capacity.

### 4. PPP inhibition induces pro-inflammatory polarization, actin reorganization, and metabolic activation in macrophages

We next examined effects induced both by pharmacological and shRNA mediated inhibition of the PPP on macrophages polarization and function.

To test whether PPP inhibition alters macrophage differentiation and activation, we assessed expression of makers delineating TAM, anti-inflammatory and tissue-regenerative subtype (M2-like), and pro-inflammatory and phagocytic-active subtype (M1-like) by flow cytometry (Fig.3A-B). We observed a trend of increased M1-like marker expression and decreased M2-like and TAM marker expression under PPP inhibition and in the PPP knockdown macrophages.

**Fig.3.**
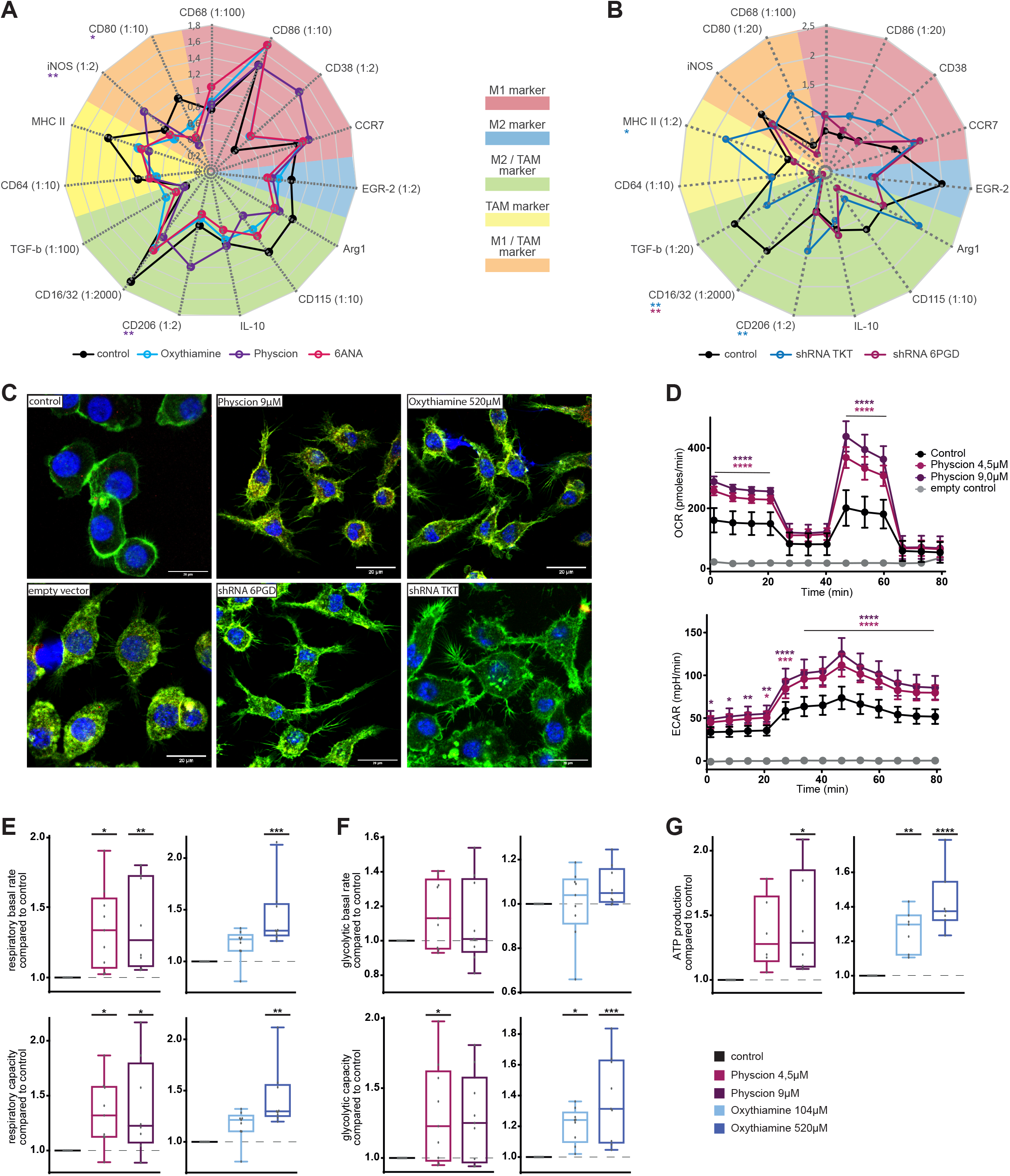
PPP inhibition improves macrophage anti-lymphoma function via pro-inflammatory polarization, actin reorganization, and metabolic activation. **(A-B)** Radar plot of surface marker expression of J774A.1 macrophages. Expression of characteristic surface marker for different macrophage subtypes measured by immune fluorescent staining. **A** compound mediated PPP inhibition, **B** shRNA mediated PPP knockdown. **(C)** Immune fluorescent microscopy of J774A.1 macrophages under compound mediated PPP inhibition and shRNA mediated PPP knockdown. Blue: Phalloidin staining of nucleus, Green: actin staining of cytoskeleton. **(D-G)** Measurement of metabolic activity of J774A.1 macrophages under compound mediated PPP inhibition by SeaHorse analysis. Inhibition of non-oxidative part of PPP by oxythiamine, inhibition of oxidative part of PPP by physcion. **D** one representative example of MitoStress test measurement of ECAR und OCR, **E** respiratory basal rate and capacity. **F** glycolytic basal rate and capacity. **G** ATP production. In **A-B** data are shown as mean of four replicates, n=4. **C** n=3. In **D** data are shown as mean of six replicates in one experiment ± SD, n=6. In **E-G** data are shown as mean ± 5-95 percentile, n=6-9. *P* values were calculated using one-way ANOVA, F using two-way ANOVA. * *p* < 0.05; \*\**p* < 0.01; \*\*\**p* < 0.001; \*\*\*\**p* < 0.0001.

To evaluate macrophage morphology, we stained actin filaments and performed fluorescent microscopy (Fig.3C). Under PPP inhibition the macrophages underwent a profound change from a round, centred appearance to a spread and outlaying phenotype with filopodia surrounding the cell body.

The metabolic status of macrophages greatly influences their activity and polarization. We thus next inhibited the PPP and assessed the glycolytic and mitochondrial activity of macrophages with the Seahorse XF Mito Stress test (Fig.3D-G, Suppl. Fig.4). We observed a general significant increase of the oxygen consumption rate (OCR) (*P*<0.0001) and extracellular acidification rate (ECAR) (*P*<0.0001) indicating an increased mitochondrial respiration and glycolytic activity and thus an increased metabolic activity of macrophages (Fig.3D). Further analysis identified significant increase of the mitochondrial basal activity (physcion *P*<0.01; oxythiamine *P*<0.001), mitochondrial maximal capacity (physcion *P*<0.05; oxythiamine *P*<0.01), glycolytic maximal capacity (physcion *P*<0.05; oxythiamine *P*<0.001), and ATP production (physcion *P*<0.05; oxythiamine *P*<0.0001) of macrophages (Fig.3E-G).

Taken together, these data show an activation of macrophages by shifted polarization, cytoskeletal reorganization, and increased metabolic activity under PPP inhibition in macrophages supporting the notion that this functionally leads to increased phagocytosis.

### 5. PPP inhibition changes the proteomic profile of macrophages towards pro-inflammatory activity

To further investigate the mediators of increased phagocytic capacity under PPP inhibition in macrophages, we performed a multi-omics (proteomic -, phosphoproteomic -, and metabolomics) screening under chemical or shRNA mediated PPP inhibition.

A uniform pattern of protein regulation by use of different independent inhibitors and PPP enzyme knockdowns was observed (Fig.4A-B). We focused on proteins involved in macrophage polarization and activation. Under compound mediated PPP inhibition, the anti-inflammatory proteins Ptgs1, Sqstm1, and Ybx3 [16][17][18] and the TAM- and M2-typicalproteins Haghand Ezr[19][20] were significantly downregulated, while the pro-inflammatory proteins Pam16 and Gosr1 were significantly upregulated (Fig.4A).

**Fig.4.**
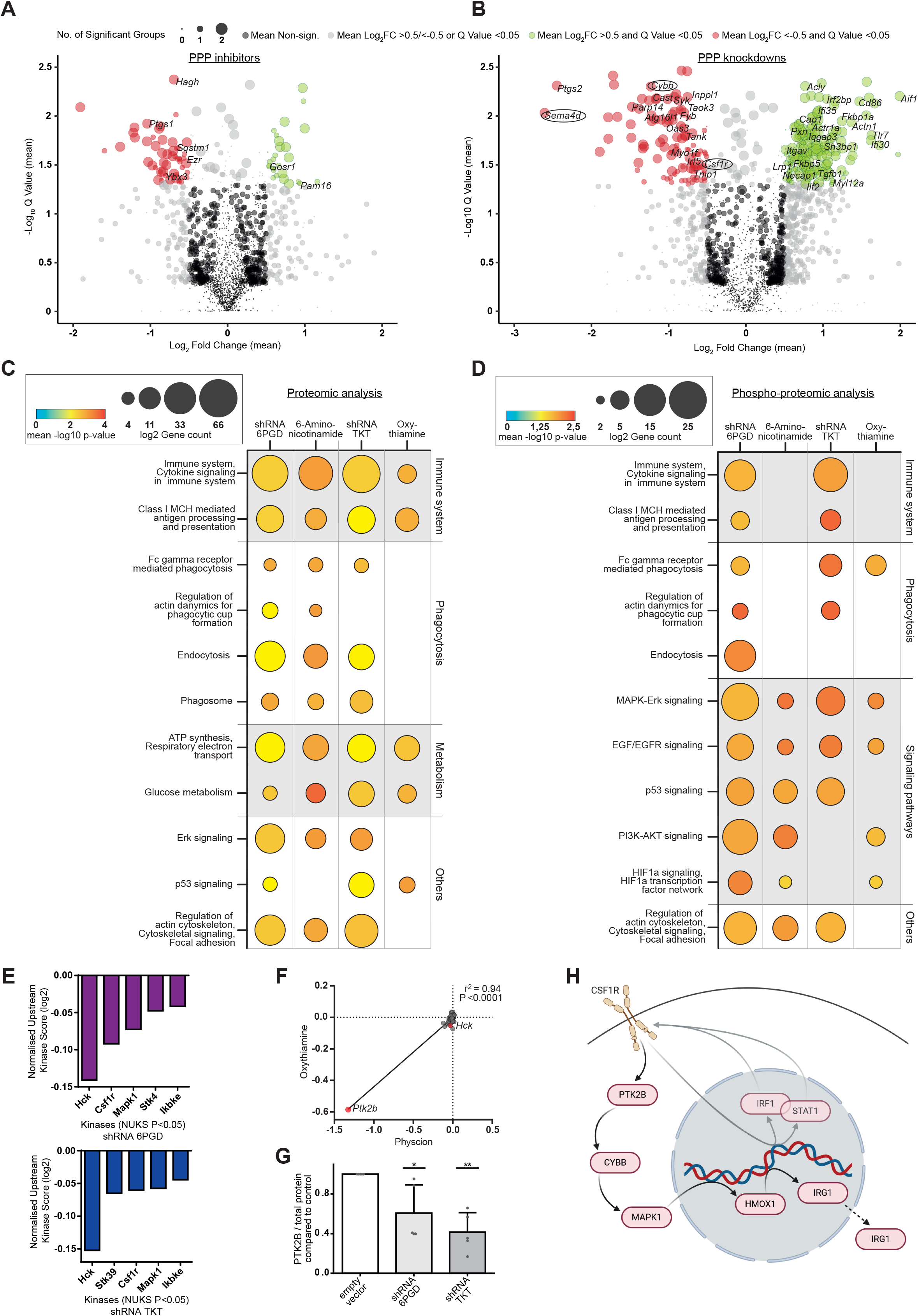
PPP inhibition changes the proteomic profile of macrophages towards pro-inflammatory activity. **(A-B)** Volcano plots showing mean change of proteomic transcription under **A** compound mediated PPP inhibition by 6-aminonicotinamide and oxythiamine compared to untreated J774A.1 macrophages, **B** shRNA mediated PPP knockdown of 6Pgd and Tkt compared to empty vector control J774A.1 macrophages. Circle size represents number of significantly changed conditions. Red circles: significantly downregulated abundance; Green circles: significantly upregulated abundance. Proteins known to participate in immune system are annotated in significant groups. **(C-D)** Pathway enrichment analysis of **C** proteomics, **D** phosphoproteomics of J774A.1 macrophages under compound mediated PPP inhibition and shRNA mediated knockdown. Protein count in listed pathways represented in circle size, -log10 *p*-value represented in heat map analysis. **(E-F)** Analysis of significant changed protein activity in *Normalized upstream-kinase score* (*NUKS*). **E** NUKS under shRNA mediated PPP knockdown of 6Pgd and Tkt, **F** integrative analysis of compound mediated PPP inhibition by physcion and oxythiamine. **(G)** Western blot analysis of Ptk2b expression in J774A.1 macrophages under shRNA mediated PPP knockdown of 6Pgd and Tkt compared to empty vector control. **(H)** Scheme of hypothesized mechanism leading to pro-inflammatory phenotype of macrophages. In **A-F** 3 biological replicates. In **G** data are shown as mean ± SEM, n=5. *P* values in **G** were calculated using one-way ANOVA. \**p* < 0.05; \*\**p* < 0.01; \*\*\**p* < 0.001; \*\*\*\**p* < 0.0001.

By using 6Pgd- or Tkt-specific shRNA an even more pronounced regulation of pro- and ant inflammatory protein expression was seen. Negative regulators of cytokine expression and pro inflammatory signalling were significantly down while promoters of pro-inflammatory activation were significantly up-regulated (Atg16l1, Cast, Csf1r, Cybb, Inppl1, Oas3, Parp14, Fkbp5, Ilf2, Tlr7) [21][22][23]. Moreover, there was a significant increase in protein expression needed for phagocytosis and phagocytic cup formation (Actn1, Actr1a, Iqgap3, Itgav, Lrp1, Myl12a, Necap1, Sh3bp1) (Fig.4B). (Phosphoproteomic analysis see Suppl. Table3)

We performed pathway enrichment analysis for functional annotation (Fig.4C-D). Similar enrichment clusters were seen by both compound mediated and shRNA mediated inhibition of the oxidative and the non-oxidative part of the PPP. A strong enrichment was seen for immune activity in all treatments (Fig.4C) including cytokine signalling, antigen processing, and antigen presentation with up to 124 involved proteins and significantly changed phosphorylation patterns in context of shRNA treatment (Fig.4D). Moreover, enrichment in proteins relevant to phagocytosis and cytoskeletal organization was observed. In line with our metabolic flux analysis (Fig.3D-G), we observed great enrichment for proteins influencing mitochondrial and glycolytic activity in all treatment conditions (Fig.4C). Particularly pathway enrichment analysis of phosphoproteomics uncovered a significant enrichment in signalling pathways important for immune signalling (Mapk-Erk, Egf/Egfr, p53, Pi3k Akt) and metabolic regulation (Pi3k-Akt, Hif1a) in PPP inhibited macrophages (Fig.4D).

To further analyse the impact of altered protein phosphorylation, we performed an adapted upstream kinase analysis on basis of Integrative Inferred Kinase Activity (INKA) analysis [24]. Protein expression, protein phosphorylation, and kinase substrate abundance were taken into account. The five most inactivated kinases are displayed (Fig.4E). Under PPP inhibition by shRNAs, a strong decrease of Hck was most prominent in the normalized upstream kinase score (NUKS) (Fig.4E). Hck supports anti-inflammatory and tissue-regenerative subtype (M2-like) polarization, TAM activity, tumor growth, and tumor cell evasion [25][26]. Furthermore, Hck activates the Csf1-receptor (Csf1r) [27]. Csf1r signalling likewise induces macrophage polarization toward a M2-like phenotype [28]. Csf1r was also one of the five most inactivated kinases in the NUKS analysis under shRNA mediated PPP inhibition (Fig.4E). Moreover, Mapk1, a downstream kinase of Csf1r signalling, was decreased (Fig.4E). In combined NUKS analysis of PPP inhibitors the Csf1r downstream kinase Ptk2b (Pyk2) was the most negatively regulated kinase (Fig.4F). Accordingly, a significant downregulation of Ptk2b under shRNA mediated PPP inhibition was detected by western blot analysis (Fig.4G, Suppl. Fig.5A). Furthermore, the most downregulated protein in both shRNA mediated knockdown macrophages, was Sema4d (Fig.4B), which is an activator of the Ptk2b pathway [29].

Following the Csf1r pathway further downstream (Fig.4H), decreased immune-regulatory gene 1 (Irg1) expression, a major node in immunosuppressive regulation of macrophages, was seen in proteomic analysis (Suppl. Table2). Changed Irg1 expression is one possible mechanism leading to altered macrophage activity and phagocytosis [30]. With the exception of Hmox1, all included signal molecules of the regarded Csf1r pathway were significantly downregulated in proteomic analysis (Fig.4H) (Suppl. Table2).

### 6. PPP inhibition modulates glycogen metabolism and the immune response signalling axis UDPG Stat1-Irg1-Itaconate of macrophages

Taking into account the critical role of Irg1 on macrophage polarization, we aimed to explore the connection between metabolic modulation, Irg1 regulation and the resulting macrophage phenotype.

PPP and glycogen synthesis activity are coupled causing suppression of both pathways if one is inhibited [31]. Moreover, glycogen metabolism influences Stat1 activity [31]. Thus, we hypothesized that inhibition of PPP would lead to suppression of glycogen metabolism with subsequent decreased UDPG production and thereby to an inhibition of P2y14 expression with following decreased Stat1 activity. Decreased Stat1 activity leads to less Irf1 and thereby to a decreased Irg1 expression, which possibly leads to functional increasing macrophage activity and phagocytosis (Fig.5A) [32][33].

**Fig.5.**
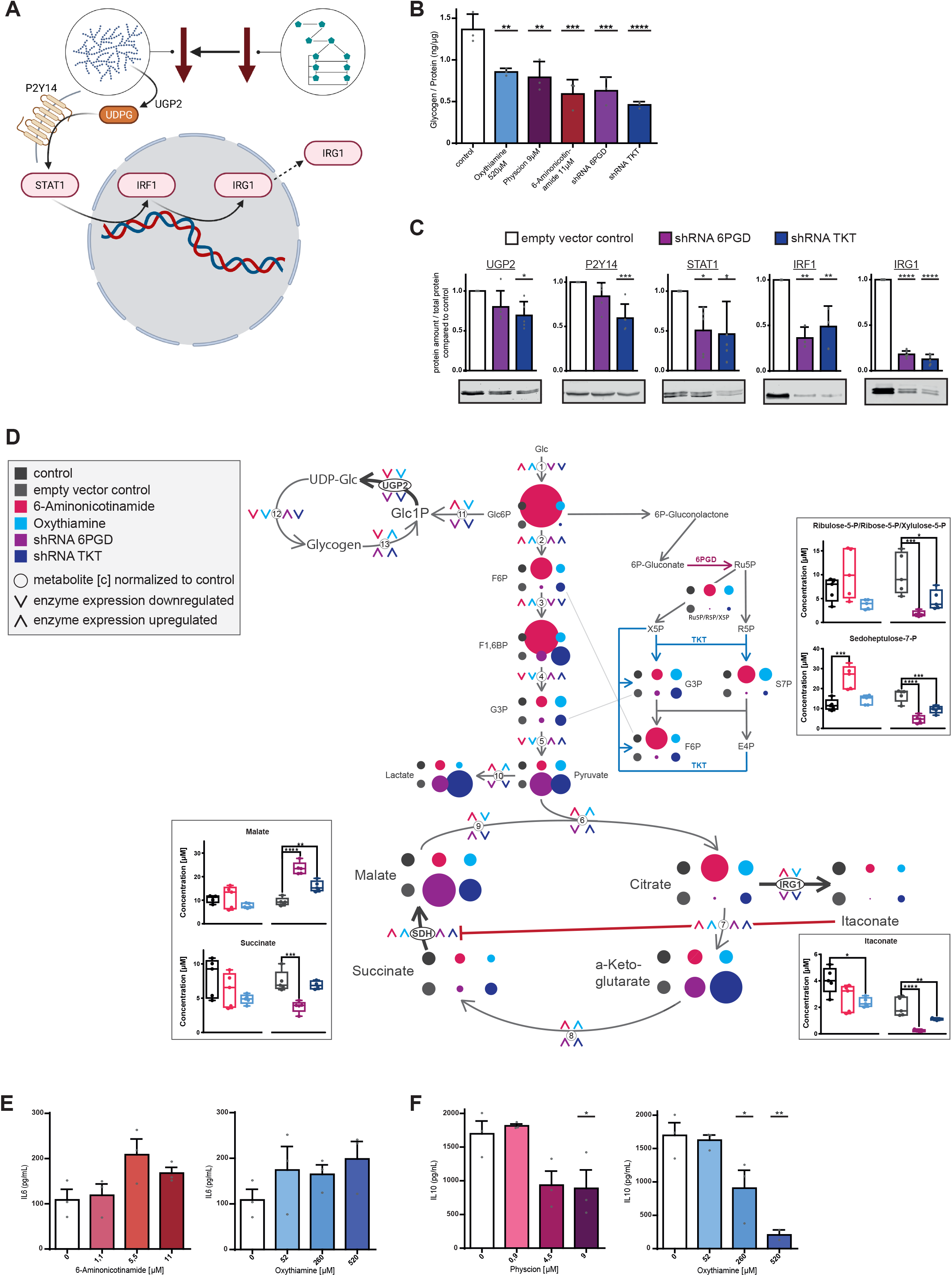
Proteomic alterations under PPP inhibition alter metabolism and immune response of macrophages. **(A)** Scheme of working hypothesis of PPP metabolism modulating immune response. **(B)** Total glycogen amount in J774A.1 macrophages under compound mediated PPP inhibition and shRNA mediated knockdown of 6Pgd and Tkt. **(C)** Western blot analysis of protein expression of hypothesized connecting pathway in J774A.1 macrophages under shRNA mediated knockdown of 6Pgd and Tkt compared to empty vector control. Mean expression displayed in bar graph analysis and one representative western blot example of each protein of interest. **(D)** Metabolomic analysis of central metabolic pathways with overlay of proteomics data under compound mediated PPP inhibition compared to untreated J774A.1 macrophages and shRNA mediated PPP knockdown of 6Pgd and Tkt compared to empty vector control J774A.1 macrophages. Relative metabolite abundance compared to respective control represented in circle size. Absolute amount of metabolites of interest displayed in bar graphs. Change in enzyme expression assessed by proteomics displayed in arrow direction. Arrow upwards: increased enzyme expression compared to respective control; arrow downwards: decreased enzyme expression compared to respective control. Inhibited enzyme reactions by compounds or shRNA mediated knockdown coloured in violet (6Pgd) and blue (Tkt). *Metabolites: E4P* erythrose-4-phosphate *, F1,6BP* fructose-1,6-bisphosphate, *F6P* fructose-6-phosphate, *G3P* glyceraldehyde-3-phosphate, *Glc* glucose, *Glc1P g* lucose-1-phosphate, *Glc6P* glucose-6-phosphate, *R5P* ribose-5-phosphate, *Ru5P* ribulose-5-phosphate, *S7P* sedoheptulose-7-phosphate, *UDP-Glc* UDP-glucose, *X5P* xylulose-5-phosphate*. Enzymes: Irg1* immune-regulatory gene 1, *Sdh* succinate dehydrogenase, *Ugp2* UDP-glucose pyrophosphorylase 2, *1)* hexokinase, *2)* glucose-6-phosphate isomerase, *3)* phosphofructokinase, *4)* aldolase, *5)* sum up of glycerinaldehyde-3-phosphate dehydrogenase, phosphoglycerate kinase, enolase, pyruvate kinase, *6)* citrate synthase, *7)* sum up of aconitase, isocitrate dehydrogenase, *8)* sum up of a-ketoglutarate dehydrogenase, succinyl-CoA synthetase, *9)* malate dehydrogenase, *10)* lactate dehydrogenase, *11)* phosphoglucomutase 1, *12)* UTP-glucose-1-phosphate uridyltransferase, *13)* glycogen phosphorylase. **(E-F)** Cytokine expression under 6Pgd inhibition by 6-aminonicotinamide or physcion and Tkt inhibition by oxythiamine in J774A.1 macrophages. **E** IL6 expression, **F** IL10 expression. **B-C** n=3-6, **D** n=1 with 3 individual replicates, **E-F** n=3. In **B-C**, **E-F** bar plots are shown as mean ± SEM, in **D** shown as Min. to Max. *P* values were calculated using one-way ANOVA. * *p* < 0.05; ** *p* < 0.01; \*\*\**p* < 0.001; \*\*\*\**p* < 0.0001.

To validate this hypothesis, we quantified glycogen levels under compound mediated and shRNA mediated PPP inhibition identifying significant decreases under all conditions (*P*<0.0001, Fig.5B). Next, we performed western blot analysis of the hypothesized pathway associated proteins identifying further significant reductions in expression (*P*<0.0001, Fig.5C) with highest decline of Irg1 amount (>80%, *P*<0.0001, Suppl. Fig.5).

As Irg1 acts as a metabolic enzyme and changes in immune activity are driven by its product itaconate, we investigated the connection between metabolism and enzyme expression by metabolomic assessment (Suppl. Table5) and overlaid the results with changes in enzyme expression of the metabolic pathways of interest detected in proteomics (Fig.5D).

Again, a uniform regulation of metabolites and enzymes was seen, especially under shRNA mediated PPP inhibition, such as significant decreased amount of exclusive 6Pgd and Tkt products ribulose-5-phosphate and sedoheptulose-7-phosphate (*P*<0.0001, Fig.5D upper right boxes). Concordant to the hypothesized coupled PPP and glycogen synthesis activity, a decrease of Ugp2 expression was seen under PPP inhibition in all conditions (upper left).

A significant decrease of the Irg1 product itaconate was observed under inhibitor mediated and shRNA mediated PPP inhibition (*P*<0.0001, Fig.5D lower right box) in line with the observed decreased Irg1 expression (Fig.5C). Itaconate is an inhibitor of succinate dehydrogenase (Sdh). Accordingly, there was an increase in Sdh expression, a significant decrease of the Sdh educt succinate (*P*<0.001) and a significant increase of the Sdh product malate (*P*<0.0001) (Fig.5D left boxes). This indicates that less itaconate production leads to less suppression of Sdh by which mitochondrial oxidative activity is increased as observed in the metabolic flux analysis (Fig.3F-G). Itaconate is known as a regulatory immunosuppressive metabolite in macrophages, which promotes anti-inflammatory IL10 secretion and inhibits pro-inflammatory IL6 secretion [30]. In line with these findings, we observed increased IL6 secretion (Fig.5E) and significantly decreased IL10 secretion (*P*<0.01, Fig.5F) by PPP inhibition.

In conclusion, we connected metabolic activity and immune regulation in macrophages via UDPG Stat1-Irg1-itaconate signalling axis provoked by PPP activity. Irg1 downregulation increases macrophage activation via itaconate reduction with subsequent metabolic activation and a pro inflammatory shift in cytokine secretion.

### 7. PPP inhibition in primary human cells increases phagocytic capacity of macrophages and decreases their bystander function

Aiming towards improved effector capabilities of macrophages in malignant disease treatment we assessed PPP inhibition in the context of human macrophages and primary leukemia co-culture systems.

To translate our findings, we isolated primary human monocytes from healthy donors and differentiated them into macrophages by M-CSF under PPP inhibition. Inhibition of both parts of the PPP significantly increased ADCP rates with more than doubled phagocytic capacity (physcion +64%; oxythiamine +110%, *P*<0.0001, Fig.6A-B). Moreover, we evaluated cytokine secretion of the human monocyte-derived macrophages post differentiation under PPP inhibition, and confirmed a similar switch toward pro-inflammatory cytokine secretion with significantly increased IL6 (*P*<0.05) and significantly decreased IL10 secretion (*P*<0.0001) (Fig.6C-D).

**Fig.6.**
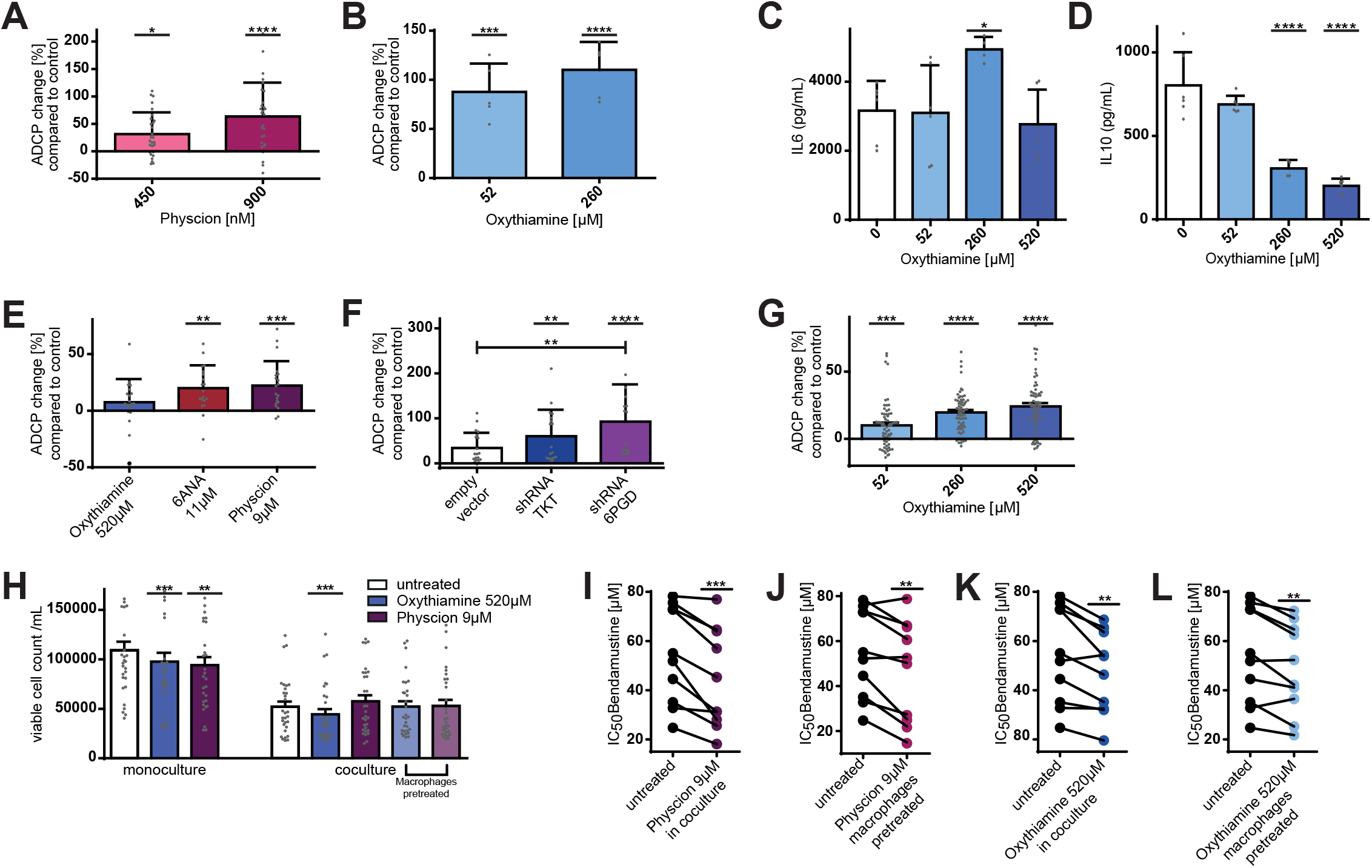
PPP inhibition in primary human cells increases phagocytic capacity of macrophages and decreases their tumor-supportive bystander function. **(A-B)** ADCP change compared to basal phagocytosis rate of human monocyte-derived macrophages. **A** ADCP change of monocyte-derived macrophages differentiated in the presence of physcion and M CSF, **B** ADCP change of monocyte-derived macrophages differentiated in the presence of oxythiamine and M-CSF. **(C-D)** Cytokine expression of monocyte-derived macrophages differentiated in the presence of oxythiamine and M-CSF. **C** IL6 expression, **D** IL10 expression. **(E-F)** ADCP change of primary CLL patient cells compared to basal phagocytosis rate of J774A.1 macrophages. **E** ADCP change under compound mediated PPP inhibition, **F** ADCP change under shRNA mediated PPP knockdown. **(G)** ADCP change of primary CLL patient cells compared to basal phagocytosis rate of monocyte-derived macrophages differentiated in the presence of oxythiamine and M-CSF. **(H)** Viability of primary CLL patient cells after incubation with PPP inhibitors physcion and oxythiamine in mono-culture and in co-culture with J774A.1 macrophages. In co-culture setting, cells were treated in parallel or macrophages were pre-treated before onset of co-culture. **(I-L)** Dose-response curve (IC_50_) for individual primary CLL patient cell samples to bendamustine treatment. Cells were incubated with bendamustine after protective macrophage co-culture with untreated J774A.1 macrophages vs. PPP inhibition. **I-J** Inhibition of 6Pgd in oxidative part of PPP by physcion, **I** co-culture treatment, **J** macrophage pre-treatment. **K-L** Inhibition of Tkt in non-oxidative part of PPP by oxythiamine, **K** co culture treatment, **L** macrophage pre-treatment. **A** n=5, **B-D, H** n=6, **E-F** n=4, **G** n=13, **I-L** n=9. In **A-H** data are shown as mean ± SEM, in **I-L** as single data plots (conjugated conditions are linked). *P* values were calculated in **A-G** using one-way ANOVA, in **H** using RM one-way ANOVA, in **I-L** using Paired t-test. \**p* < 0.05; \*\**p* < 0.01; \*\*\**p* < 0.001; \*\*\*\**p* < 0.0001.

To address effector function of macrophages under PPP inhibition in the context of primary human leukemia cells, primary CLL patient cells were used for ADCP assays with macrophages. A significant increase of phagocytosis was observed under PPP inhibition (+22%, *P*<0.001, Fig.6E) and by using knockdown macrophages with a nearly doubled ADCP rate by using 6Pgd knockdown macrophages (Tkt +60%; 6Pgd +92%, *P*<0.0001, Fig.6F).

To evaluate phagocytosis in a fully human setting, we performed ADCP assay with primary human monocyte-derived macrophages differentiated in the presence of PPP inhibitors and primary leukemia patient cells. A significantly increased phagocytic capacity was observed (+24%, *P*<0.0001, Fig.6G).

We have demonstrated that also primary indolent lymphoma and primary human macrophages are affected by PPP.

Beyond the inefficient phagocytic function, macrophages in the TME exert direct supportive effects on tumor cells. CLL cells depend on macrophages as “nurse-like” bystander cells to survive [34]. Macrophages in the microenvironment of CLL are polarized towards tumor promoting TAMs and support CLL cells by chemokine secretion and immunosuppressive signalling. We therefore evaluated the effect of PPP-inhibited macrophages on primary CLL cells in co-culture. Interestingly, PPP inhibition in mono-cultured primary CLL cells decreased their viability significantly (*P*<0.0001, Fig.6H, left). Co-culture inhibition of the non-oxidative part of the PPP reduced the viability of primary CLL cells significantly (*P*<0.01, Fig.6H, right).

As TAMs are also important mediators in chemotherapy resistance, we evaluated if the co-cultivation under PPP inhibition might affect the susceptibility of primary CLL cells towards apoptosis by chemotherapy. Primary CLL patient cells were cultured with macrophages under PPP inhibition and subsequently exposed to the chemotherapeutic agent bendamustine. We observed significantly increased bendamustine-induced apoptosis among primary CLL cells under PPP inhibition of both parts of the PPP (*P*<0.001, Fig.6I-L). This boost in primary CLL cell apoptosis was achieved by PPP inhibition in the co-culture (*P*<0.001, Fig.6I, K) and by macrophages pre-treatment with PPP inhibitors before exposing them to primary CLL cells (*P*<0.01, Fig.6J, L).

These observations underline the role of altered macrophage support under PPP inhibition in the TME such as direct leukemia cell support or resistance to chemotherapy.

### 8. PPP inhibition increases macrophageś maturation and pro-inflammatory polarization in vivo

To evaluate if PPP inhibition preserves its effect on macrophages also *in vivo*, we treated C57BL/6J mice with the PPP inhibitor S3 [14] for 7 days and analysed macrophages of different compartments afterwards.

For investigating the myelopoiesis under PPP inhibition, we performed surface staining of bone marrow cells and observed a significant increase of cells in the LSK compartment (Lin ^-^, Sca-1^+^, c-Kit^+^) under S3 treatment (*P*<0.0001, Fig.7A). PPP inhibition significantly increased the amount of hematopoietic stem cells (HSC; *P*<0.05, Fig.7B) and multipotent progenitor (MPP) cells (Fig.7B), including MPP2, MPP3, and MPP5, which are known to fuel the myeloid compartment, especially under demand of myeloid expansion (MPP2 *P*<0.001; MPP3 *P*<0.01; MPP5 *P*<0.01, Fig.7B) [35][36]. This coincided with a significantly increased frequency of myeloid progenitor cells (*P*<0.01, Fig.7C) and macrophages (*P*<0.001, Fig.7C) in the bone marrow, indicating propelled myelopoiesis under PPP inhibition.

**Fig.7.**
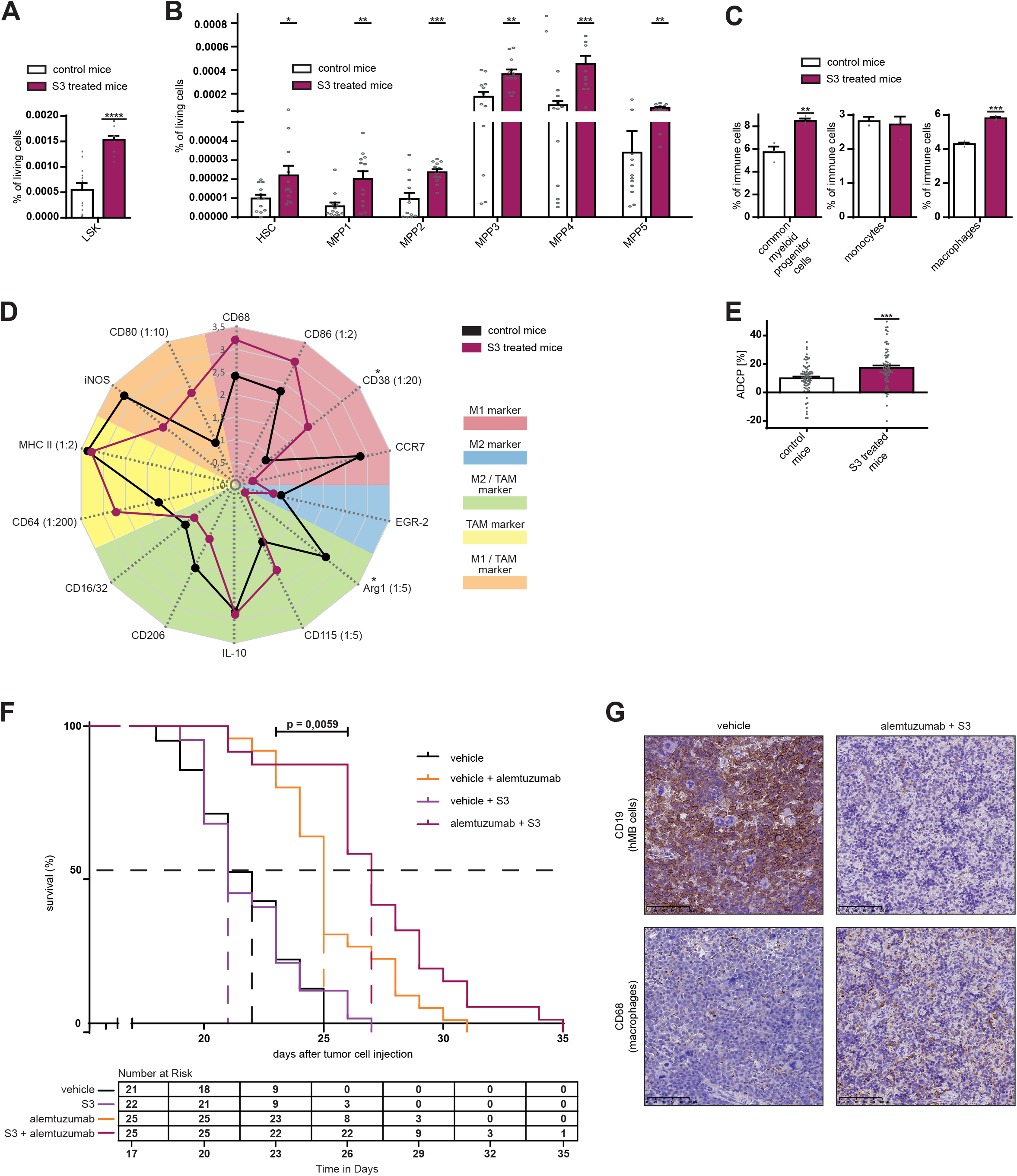
PPP inhibition increases myelopoiesis and macrophages activity *in vivo* and improves treatment response in an aggressive humanized lymphoma mouse model. **(A)** Progenitor cell compartment LSK (Lin ^-^, Sca-1 ^+^, c-Kit^+^) in bone marrow of C57BL/6J mice treated with vehicle (control) or PPP inhibitor S3 i.p. for 7 days. **(B)** Multipotent progenitor (MPP) subsets in bone marrow of C57BL/6J mice treated with vehicle (control) or PPP inhibitor S3 i.p. for 7 days. HSC (CD34^-^, CD48^-^, CD150^+^, CD135^-^), MPP1 (CD34^+^, CD48^-^, CD150^+^, CD135^-^), MPP2 (CD34^+^, CD48^+^, CD150^+^, CD135^-^), MPP3 (CD34^+^, CD48^+^, CD150^-^, CD135^-^), MPP4 (CD34^+^, CD48^+^, CD150^-^, CD135^+^), MPP5 (CD34^+^, CD48^-^, CD150^-^, CD135^-^). **(C)** Percentage of myeloid lineage cells of whole cell amount in bone marrow of NSG mice transfected with hMB cells, treated with vehicle or PPP inhibitor S3 i.p. for 12 days and euthanized on day 15. Common myeloid progenitor cells (CD41 ^+^, CD34 ^+^), monocytes (Ly6c ^+^, CX3CR1^+^), macrophages (F4/80 ^+^, CD64^+^). **(D)** Expression of characteristic surface marker for different macrophage subtypes on peritoneal macrophages measured by immune fluorescent staining. C57BL/6J mice treated with vehicle (control) or PPP inhibitor S3 i.p. for 7 days. **(E)** ADCP assay of bone marrow derived macrophages. C57BL/6J mice treated with vehicle (control) or PPP inhibitor S3 i.p. for 7 days, macrophages differentiated out of femoral bone marrow with M-CSF. **(F)** Survival curve of aggressive lymphoma (hMB) bearing mice treated with PPP inhibitor S3 +/- therapeutic antibody alemtuzumab. **(G)** One representative example of immunohistochemical staining of hMB cells (CD19 ^+^) and macrophages (CD68 ^+^) in spleen of aggressive lymphoma (hMB) bearing mice treated with vehicle or alemtuzumab + S3. **A-B** n=12, **C** n=3, **D** n=10, **E** n=10, **F** n=25. In **A-C, E** data are shown as mean ± SEM, in **D** data are shown as mean of ten replicates. *P* values were calculated in **A-E** using **Unpaired** t-test, and in **F** using Benjamini-Hochberg test. \**p* < 0.05; \*\**p* < 0.01; \*\*\**p* < 0.001; \*\*\*\**p* < 0.0001.

We further investigated the polarization of *in vivo* macrophages and detected a shift of polarization away from M2-like subtype with significantly decreased Arg1 expression (*P*<0.05), and towards the pro-phagocytic M1-like subtype with significantly increased CD38 expression (*P*<0.05) upon PPP inhibition *in vivo* (Fig.7D) (other compartments see Suppl. Fig.7A). These findings are in line with our *in vitro* observations.

Moreover, we generated bone marrow derived macrophages of S3 treated and control mice and performed ADCP assays. *In vivo* PPP inhibition significantly increased phagocytic activity of macrophages (+74%, *P*<0.001, Fig.7E) compared to bone marrow derived macrophages of controls.

Altogether, we have shown that PPP inhibition *in vivo* activates macrophages’ inflammatory polarization and maturation as well as their phagocytic capacity, which increases their anti-tumor function *in vivo*.

### 9. PPP inhibition boosts anti-leukemic treatment and thereby prolongs survival in an aggressive lymphoma mouse model

To focus on the therapeutic effect of PPP inhibition *in vivo*, we evaluated treatment effects in an aggressive lymphoma mouse model.

We used the humanized double-hit lymphoma mouse model (hMB) [37], which is amenable for modelling human specific antibody therapy, and treated the mice with the therapeutic antibody alemtuzumab and the PPP inhibitor S3. As the lymphoma reflects aggressive disease, untreated mice died rapidly after tumor cell injection (median survival (MS) 22 days, Fig.7F). By treatment with S3 only, this rapid tumor progression persisted (Fig.7F). As shown in our previous work, treatment with alemtuzumab increases survival significantly in this aggressive lymphoma mouse model [6] (MS 25 days, *P*=1.4e-05, Fig.7F, Suppl. Fig.7B). By adding the PPP inhibitor S3 to alemtuzumab an additional significant prolongation of mouse survival was achieved in comparison to antibody treatment only (MS 27 days, *P*=0.0059, Fig.7F, Suppl. Fig.7B) with a stable increased number at risk up to day 25 (survival of 88%, Fig.7F). Immunohistochemical analysis of spleens showed a marked reduction of CD19^+^ lymphoma cell infiltration with concomitant increase of CD68 ^+^ macrophage infiltration after treatment with alemtuzumab and S3 in comparison to vehicle control (Fig.7G, Suppl. Fig.7C).

We finally demonstrate *in vivo* that PPP inhibition in the context of a highly aggressive lymphoma model increases the efficacy of antibody therapy to prolong overall survival significantly.

## Discussion

Tumor associated macrophages (TAMs) are key drivers in multiple malignancies associated with poor outcome and diminished efficacy of immunotherapies [1][3][6]. The influence of glucose and mitochondrial metabolism on macrophages’ polarization and activity has been established [9][38][39][10], while the pentose phosphate pathway (PPP) has not yet been implicated in TAM regulation. Here, we show that modulation of the PPP in TAMs serves as a robust regulator of phagocytic function and macrophage activity and prolongs survival in aggressive B-cell lymphoma therapy.

Metabolic inhibition screening in lymphoma-macrophage co-cultures emphasized detrimental effects on macrophage function for the majority of investigated pathways. Only inhibition of the PPP showed a significant increase of phagocytosis with a synergistic effect on effector and target cells. We identified PPP as a robust and reproducible regulator of phagocytic activity in multiple *in vitro* and *in vivo* macrophage model systems both by compound and genetic targeting. The increased phagocytosis rate appeared also under hypoxic conditions as an approximation of physiological status of therapy-refractory niches of lymphoma – the lymph nodes and bone marrow [40][41] – indicating therapeutic efficacy by metabolic inhibition in contrast to other therapy modalities in these niches.

Previous reports demonstrated reduced cancer and leukemia cell growth in mice upon PPP inhibition [14][42]. Especially in CLL, macrophages play a pivotal role as supportive bystander cells in the TME. Without the support of TAMs, CLL cells would undergo spontaneous apoptosis [34]. We have shown here that PPP inhibition also diminishes this pro-survival bystander function of macrophages. As mono-cultured primary CLL cellś viability was reduced upon PPP inhibition, a direct negative effect on primary CLL cells’ survival is to be expected. In line with that, a higher susceptibility of primary CLL cells to chemotherapeutic treatment under PPP inhibition in macrophage co-culture was demonstrated. PPP inhibition might serve as a sensitizer to conventional chemotherapeutic regimens.

Alterations of the PPP enzymes 6PGD and TKT have been previously described in many cancer types [43][44][45][46][47][48][49]. Overexpression of TKT was closely associated with aggressive hepatocellular carcinoma features [50], and 6PGD was shown to promote metastasis [51] and cisplatin resistance [52]. Suppression of 6Pgd attenuates cell proliferation and tumor growth [42], overcomes cisplatin resistance [52] and Tkt inhibition sensitizes cancer cells to targeted therapy [50]. PPP inhibition by physcion, S3 or 6ANA has demonstrated anti-tumorigenic effects in leukemia, glioblastoma, and lung cancer cells [53][54][55], as well as in chemotherapeutic resistant AML cells [49], without affecting proliferation of non-malignant cells [42]. Knockdown of Tkt profoundly reduced growth of metastatic lesions, while Tkt inhibition in normal cells did not have suppressive effects [50]. Beside its effect on cancer cells, it was investigated that 6Pgd inhibition in CD8 ^+^ T-cells led to an increased effector function with higher tumoricidal activity [56].

We have previously shown that macrophage effector polarization is crucial in therapeutic antibody based regimens of B-cell lymphoma and can be modulated by chemotherapeutic agents, targeted drugs or extracellular vesicle mediated cell-cell communications [57][58][59]. We now identified macrophage metabolism as a crucial switch of macrophage effector function in lymphoma.

We demonstrated that increase of phagocytosis is driven directly by PPP enzyme inhibition, rather than metabolite accumulation as indicated by the response pattern to PPP intermediate supplementation to macrophages. Non-exclusive PPP metabolites did not influence phagocytosis, possibly due to degradation via glycolysis (G3P, F6P) or nucleotide synthesis (R5P) before entering PPP flux. In contrast, exclusive PPP metabolites altered phagocytosis activity: supplementation of the G6pd product 6-phosphogluconate, the 6Pgd product Ru5P, and the Tkt product S7P increased phagocytosis, while supplementation of the Tkt educt E4P decreased phagocytic activity in macrophages. This points to a feedback inhibition of the respective PPP enzymes and emphasizes the enzyme inhibition as driving force for increased phagocytosis.

PPP is a central linker between glucose metabolism, amino acid biosynthesis, fatty acid metabolism and redox homeostasis [43]. A gain in metabolic activity was observed under PPP inhibition with increased glycolytic and mitochondrial capacity causing enhanced ATP production, thereby fuelling macrophages’ activation. An increase of glycolysis is well described within the phenotypical switch to pro-inflammatory macrophages [60]. Furthermore, we observed changed cytoskeletal organization under PPP inhibition. The enlarged cell size and widespread cytosol protrusions reflect an activated state of macrophages and point to a higher disposition for phagocytosis, as filopodia initiate phagocytosis by approaching the target prior engulfment. Assessment of macrophage polarity demonstrated a decrease of marker associated with M2-like macrophages and TAM, which represent immunosuppressive and tumor-promoting macrophage subtypes [3][61][62], while exclusive M1 marker, expressed on pro-inflammatory macrophages, were increased [63][64][65][66].

In total, the restriction of one metabolic pathway – the PPP – gives rise to numerous paths of activation which renders a profound alteration of phenotype and particular phagocytic activity in macrophages. Thereby the anti-tumor function could be improved from independent directions.

Our detailed multi-omics and functional analysis provides evidence that these effects are directly related to 6Pgd and Tkt enzyme activity loss, which polarizes macrophages to a pro-inflammatory phenotype through downregulation of Stat1 and Irg1. The functional switch between PPP enzyme activity and subsequent polarization program are Csf1r expression or activity of glycogen synthesis. Csf1r activation induces a signalling cascade, which leads to Hmox-1 expression [67][68]. Hmox-1 in turn induces Irg1 expression [69], a central inhibitory regulator of macrophage activation [30]. With exception of Hmox-1, all participants of this Csf1r pathway were shown to be downregulated significantly in our proteomic screening. Via Csf1r signalling, macrophages are polarized toward a M2-like or TAM phenotype by also direct activation of Erk1/2 (Mapk1/2) and Hck signalling [26][27][28][70][71][72]. It has been shown that inhibition of Hck activity reduces the amount of

TAMs, colon cancer growth and tumor cell invasion [25]. Both, Mapk1 and Hck activity was shown to be decreased under PPP inhibition in upstream kinase analysis.

Considering the relevant role of Csf1r in macrophage ontogeny, activation, and polarization, reduced Csf1r expression might be responsible for macrophage activation under PPP inhibition. As inhibition of Csf1r is tolerable in adult mice [73], Csf1r blockade might be a promising strategy to increase macrophages activity in context of tumor therapy. Several CSF1/CSF1R inhibitors are currently under clinical investigation [74]. Nevertheless, as CSF1R is a macrophage exclusive receptor, only a macrophage exclusive effect could be achieved by using CSF1R inhibition, in contrast to the above described multi-cellular effects of 6PGD and TKT inhibition.

As Csf1r signals via Stat1 [75][76], the observed reduction of Stat1 expression might be due to reduced Csf1r expression and a putative link between metabolism and immune regulation. Recently, PPP inhibition was linked to inhibition of glycogen synthesis causing decreased UDPG production [31]. UDPG is a signalling molecule, which activates the P2y14 receptor (P2y14r). P2y14r activation in turn increases Stat1 expression and Mapk1 phosphorylation [77]. Considering that Stat1 is a major regulator of Irf1 expression [32], which is the transcription factor of Irg1 [78], an impact of PPP inhibition on Irg1 expression is likely. Irg1 is almost exclusively expressed in activated immune cells and a key driver of immune inhibition *via* itaconate production [30]. Itaconate inhibits glycolysis and mitochondrial activity by Sdh inhibition [79][80][81] and promotes anti-inflammatory macrophage phenotype [82] and tumor growth by increased ROS secretion by TAMs [83]. In line with this hypothesis, we observed decreased expression of all UDPG-Stat1-Irg1-itaconate pathway proteins under PPP inhibition, particularly lower amounts of itaconate and an increased Sdh activity. This subsequently resulted in pro-inflammatory cytokine switch, phenotypic shift towards M1-like macrophages and a diminished primary leukemia cell support. The metabolic alterations observed here, support the hypothesis of the connection between PPP inhibition, itaconate abundance and immune regulation.

The transcription factor Irf1 also induces iNos expression [78], consistent with the observed decreased iNos expression under PPP inhibition. High iNos expression and activity has been correlated with malignancy and poor survival in several solid tumors and leukemia [84].

In turn, P2y14r expression in T-lymphocytes was shown to inhibit T-cell proliferation and activation [85], also suggesting PPP inhibition to be a regulator of adaptive immune responses.

As reduced expression of Irf1 by PPP inhibition also influences Stat1 and Csf1r expression [86], Irf1 appears as the junction between all recapitulated pathways, which are leading to the changed macrophage activity and polarization.

As we initially intended to utilize metabolic modulation to improve macrophage function in human therapy regimens, we evaluated treatment efficacy also using CLL patient cells and primary human macrophages. We could demonstrate here a highly significant increase of phagocytosis in primary human cells, indicating efficacy for potential clinical use.

To model lymphoma in human patients, we performed *in vivo* experiments. Mice were injected with humanized double-hit lymphoma cells (hMB) and treated with the CD52-antibody alemtuzumab and the PPP inhibitor S3, whose low toxicity and high effectiveness in treatment of other tumor entities was demonstrated before [14][42]. In this aggressive lymphoma mouse model [37], PPP inhibition has an additional effect on antibody therapy leading to significant prolonged overall survival in comparison to antibody treatment only with associated increased macrophage lymphoma infiltration. PPP inhibition *in vivo* increased myelopoiesis and gave rise to progenitor cell expansion, indicating increased provision of a variety of immune effector cells. Macrophages displayed pro inflammatory polarization and significantly increased phagocytic capacity after PPP inhibition *in vivo*. Here we have proven *in vivo* efficacy of PPP inhibition leading to macrophage activation and improving therapy response by antibody-mediated phagocytic clearance of lymphoma with prolongation of overall survival.

In conclusion, PPP inhibition serves as immune modulatory therapy repolarizing macrophages. We indicate metabolic modulation as a key mechanism of macrophage regulation. PPP inhibition causes diminished Irg1 expression leading to reduced anti-inflammatory itaconate production. Our work indicates PPP inhibition as a dual principle therapy targeting cancer cells [42][14] and their immune microenvironment simultaneously, especially in context of antibody-based regimens.

## Acknowledgements

We are indebted to our patients who contributed tissue and blood samples to this study. This work was funded by the Deutsche Forschungsgemeinschaft (DFG, German Research Foundation) KFO28 and SFB1530 (SFB-Geschaeftszeichen – 455784452, Project B02). C.P.P. was supported by the “Foerderprogramm Nachwuchsforschungsgruppen NRW 2015–2021”, CAP Program of the Centre for Molecular Medicine Cologne and a research grant by Gilead Sciences. A.C.B. was supported by „Studentische Forschungsfoerderung/ Begabtenfoerderung” of Koeln Fortune program of the medical faculty of University of Cologne. We would like to thank Thomas Wunderlich for advice and critical reading. We are grateful for technical assistance from the CECAD Imaging and animal facilities.

## Author Contribution

Study Design: A.C.B., N.N. and C.P.P.; Data Analysis and Acquisition: A.C.B., S.J.B., M.K., S.C., M.M., R.B., H.H.B., D.V., J.L.N., R.L., C.R.P., J.S., A.V., Bioinformatic Analyses: A.C.B., E.I., S.J.B., J.L.N., C.B. and M.K. Analytical tools: H.W., M.K. and C.B. Clinical Samples and Annotation: C.P.P. and M.H.; Study Supervision and Funding: C.P.P.; Manuscript Preparation: A.C.B., S.J.B. and C.P.P.

## Declaration of Interests

The authors declare no competing interests.

## Inclusion and diversity

We worked to ensure gender balance in the recruitment of human subjects. We worked to ensure sex balance in the selection of non-human subjects. We worked to ensure diversity in experimental samples through the selection of the cell lines. One or more of the authors of this paper self identifies as an underrepresented ethnic minority in their field of research or within their geographical location. One or more of the authors of this paper self-identifies as living with a disability. While citing references scientifically relevant for this work, we also actively worked to promote gender balance in our reference list. We avoided “helicopter science” practices by including the participating local contributors from the region where we conducted the research as authors on the paper.

## Notes

### Competing Interest Statement

The authors have declared no competing interest.

