## Supplementary material for "Pentose Phosphate Pathway Inhibition activates Macrophages towards phagocytic Lymphoma Cell Clearance": Methods

**STAR METHODS**

**RESOURCE AVAILABILITY**

**Lead contact**

**Materials availability**

Oligonucleotides in this study have been generated by SIGMA ALDRICH, catalogue number is available in this paper´s supplemental information.

**Data and code availability**

Proteomic data have been deposited at PRIDE database (**Project Name:** Macrophage regulation by the Pentose Phosphate Pathway in the TME, **Project accession:** PXD042428; Reviewer account details: Username:, Password: SMwjxsiL), and are publicly available as of the date of publication. Accession numbers are listed in the key resources table.

Phosphoproteomic data have been deposited at PRIDE database (**Project Name:** Macrophage regulation by the Pentose Phosphate Pathway in the TME, **Project accession:** PXD042428; Reviewer account details: Username:, Password: SMwjxsiL) and are publicly available as of the date of publication. Accession numbers are listed in the key resources table.

All original code is available in this paper´s supplemental information.

Any additional information required to reanalyse the data reported in this paper is available from the lead contact upon request.

**EXPERIMENTAL MODEL AND SUBJECT DETAILS**

**Mouse strains**

Wild-type NOD.Cg-Prkdcscid Il2rgtm1Wjl/SzJ (NSG) and C57BL/6 mice were from Jackson laboratory (ME, USA). NSG is an immune-deficient strain with compromised generation of lymphocytes, natural killer cells, macrophages, and immunoglobulins caused by lacking expression of PRKDC and the γ-chain of interleukin-2 receptor. C57BL/6 is a commonly used immune competent inbreeding strain. Animals (female and male; 0-40 weeks) were maintained under specific pathogen free conditions in line with European Union regulations. Experiments were approved by local ethical review (LANUV (Landesamt für Natur, Umwelt und Verbraucherschutz Nordrhein-Westfalen)) and were carried out under the authority of Michael Michalik M. Sc., (Uniklinik Köln, CECAD Research Centre, Ebene 2.2 – AG Pallasch, Joseph-Stelzmann-Str. 26, 50931 Köln) project license.

Mice were kept under cage conditions at 20-22 °C. Littermates of the same sex were randomly assigned to experimental groups. Adult mouse weight is around 23,4 grams.

**Cell lines**

HEK293T-CAF40-null cells were from DSMZ, hMB cells were generated by Leskov et al. [1], J774A.1 macrophages were from ATCC, THP1 monocytes were from DSMZ. All cell lines were cultured on 6 well plates or 10cm dishes from Corning and incubated with 5% CO_2_ at 37,0 °C. J774A.1 cells, hMB cells, and HEK293T cells were cultured in DMEM containing 10% fetal bovine serum and 1% penicillin/streptomycin. THP1 cells were cultured in RPMI 1640 medium containing 10% FBS and 1% penicillin/streptomycin.

**Primary cells**

Primary CLL patient cells and primary monocytes from healthy donors were collected from buffy coats donated by blood bank of University Hospital of Cologne. Cells were cultured on 6 well plates or 10cm dishes from Corning and incubated with 5% CO_2_ at 37,0 °C. The cells were cultured in RPMI 1640 medium containing 10% FBS and 1% penicillin/streptomycin.

Monocytes of buffy coats were separated by CD14 anti-human magnetically labelled MicroBeads from Miltenyi Biotec.

**Microbe strains**

5-alpha competent E. coli from New England Biolabs Inc. were used for plasmid generation.

**METHOD DETAILS**

**Antibody-dependent cellular phagocytosis assay (ADCP)**

1x10^4^ J774A.1 cells were plated out in 100µL media per well in a 96 well plate. After 24hrs of attachment hMB cells were added in a macrophage : hMB ratio 1 : 15. Compounds were added in increasing concentration up to maximal non-toxic concentration. Alemtuzumab was added to every second well in a concentration of 10µg/mL. Wells were filled up with macrophage medium up to volume of 250µL. After 18hrs of incubation remaining hMB cells per well (GFP positive) were measured using Miltenyi MacsQuant VYB flow cytometer. Out of the absolute GFP positive cell count the antibody-dependent cellular phagocytosis rate was calculated with the formula 100-(100*(total GFP^+^ antibody-treated well/total GFP^+^ antibody-untreated well)). The calculated ADCP rate was compared to basal ADCP rate of untreated control cells (set as 100%) by division of the calculated ADCP rates (= ADCP change). For ADCP assays performed with THP1 cells, the amounts were subtracted to avoid bias as basal phagocytosis rate of THP1 cells is low (= ADCP difference).

For pre-treatment assays macrophages respectively hMB cells were treated with increasing concentration of the compounds up to maximal non-toxic concentration. After 24hrs of incubation the cells were washed three times and the assay was performed as described above without addition of the compounds to co-culture.

Performing the assay with THP1 cells, the antibody obinutuzumab was used in a concentration of 1µg/mL.

Compounds and antibodies were diluted in media of used macrophage type respectively in hMB medium in case of hMB pre-treatment ADCP assays.

For ADCPs in hypoxia, cells were incubated under hypoxic conditions with 1,5% O_2_ and 5% CO_2_.

For ADCPs with CLL patient cells, CLL patient cells were used instead of hMB cells.

For ADCPs with primary human macrophages, 2x10^4^ primary human macrophages and 3x10^5^ hMB cells were used per well. As antibody daratumumab in a concentration of 10µg/mL was used.

For ADCPs with primary murine macrophages, 5x10^4^ primary murine macrophages and 1,5x10^5^ hMB cells were used per well. As antibody daratumumab in a concentration of 10µg/mL was used.

**Antibody-independent cellular phagocytosis assay (AiDCP)**

Experiment was performed like ADCP without the addition of antibody. Remaining hMB cells were compared to hMB mono-culture cell count under inhibitor treatment. After 18hrs of incubation remaining hMB cells per well were measured using Miltenyi MacsQuant VYB flow cytometer. Out of the absolute GFP positive cell count the antibody-independent cellular phagocytosis rate was calculated with the formula 100-(100*(total GFP^+^ macrophage co-culture well/total GFP^+^ hMB mono-culture well)).

**Bone marrow derived macrophage generation**

Femur were flushed with DMEM media. Erythrocytes were lysed by addition of 2mL ACK lysis buffer to cells. Reaction was stopped by addition of 50mL cold PBS. Cells were re-suspended in media and plated out on 10cm cell culture plates for 24hrs. Non adherent cells were collected and plated out in a concentration of 6x10^5 cells /mL on 10cm cell culture plates in media and 15% feeder media 1 and 2. Feeder media was collected from L-929 cells after incubation with RPMI media for one week (feeder media 1) and three weeks (feeder media 2). On day three additional 4mL media with 15% feeder media 1 and 2 and 50ng murine recombinant M-CSF were added. On day seven media was replaced by 10mL media. On day eight the plates were washed two times and adherent macrophages were detached by scraping. Macrophages were plated out for further experiments and were incubated for 24hrs for recovering before further experiments were performed.

**CLL patient cell co-culture**

5x10^4^ J774A.1 cells were plated out in 1mL media on a 24 well plate and were incubated for 5hrs. Three samples per condition were treated with the PPP inhibitors for macrophages pre-treatment. Cells were incubated for 24hrs. Pre-treated cells were washed three times. Viability of CLL patient cells was measured by 7AAD plus AnnexinV staining. Therefor, 2µL 7AAD stain, 2µL AnnexinV stain and 46µL 1% ABB were added to washed cells, cells were incubated for 20min at 4°C and additionally 50µL 1% ABB was added. Readout was performed immediately using Miltenyi MacsQuant X flow cytometer. 7,5x10^5^ viable CLL patient cells in 1mL media were added to the macrophages and CLL mono-culture wells were plated out under addition of 1mL DMEM. Three samples per condition were treated with PPP inhibitors for co-culture treatment. The co-culture was incubated for three days. CLL cells were re-suspended by pipetting and supernatant was transferred into eppies. On U-bottom plates 7AAD plus AnnexinV staining (see above) were performed and cell count was measured using Miltenyi MacsQuant X flow cytometer.

**CLL patient cell chemotoxicity stain**

CLL patient cells after CLL co-culture performance were transferred into 96 U-bottom plates. 25µL of each Bendamustine concentration was added to one well of each condition. Cells were incubated for 48hrs. Cells were washed and 7AAD plus AnnexinV staining was performed (see above).

Chemotoxicity assay was also performed by adding the cells to 1x10^4^ J774A.1 cells per well plated out the day before. After co-incubation for 48hrs, wells were mixed up and supernatant was transferred to a 96 U-bottom plate to perform 7AAD plus AnnexinV staining of the CLL patient cells (see above).

**ELISA**

7x10^5^ J774A.1 cells in 2mL media per well were plated out on a 12 well plate and incubated for 24hrs. Each inhibitor was added to two wells and cells were incubated for 24hrs. 100ng/mL LPS was added to one well of each condition. After incubation of 16hrs supernatant was transferred into Eppendorf tubes. The supernatant was centrifuged at 300g for 5min and transferred to new Eppendorf tubes. BioLegend ELISA kit for TNF-α, IL10, and IL6 was used and protocol performed as from BioLegend mentioned. Readout was performed by fluorescence intensity measurement using FLUOStar OPTIMA.

**Immune fluorescent Microscopy**

Microscopy coverslips were washed three times in ethanol and autoclaved. Two microscopy coverslips per well were placed on a 6 well plate. 1x10^6^ J774A.1 cells in 1mL media per well were plated out, treated with the PPP inhibitors and incubated for 24hrs. Cells were washed with 5mL DPBS for 5min. 2mL PFA was put on the cells and incubated for 10min. Cells were washed three times with 5mL DPBS for 5min. Cells were incubated with 2mL of DPBS with 0,25% Triton X for 3min. Cells were incubated with 2mL of DPBS with 0,125% Triton X plus 5% BSA for 1hr. Coverslips were taken off the 12 well plate and dried on a paper towel. Mitochondrial antibody TOM20 F-10 was diluted 1:500 in DPBS plus 0,125% Triton X plus 2,5% BSA, 100µL was pipetted on the coverslips and they were incubated for 2hrs. Coverslips were washed three times with 100µL DPBS with 0,125% Triton X. 2^nd^ mitochondrial antibody (Alexa Fluor 647 α mouse) was diluted 1:1000 in DPBS plus 0,125% Triton X plus 2,5% BSA, 100µL was pipetted on the coverslips and they were incubated for 1hr. Coverslips were washed three times with 100µL DPBS with 0,125% Triton X. Actin antibody (Phalloidin Alexa Fluor568) was diluted 1:1000 in DPBS with 0,125% Triton X plus 2,5% BSA, 100µL was pipetted on the coverslips and they were incubated for 1hr. Coverslips were washed nine times with 100µL DPBS with 0,125% Triton X. Nuclear antibody (DAPI) was diluted 1:10000 in DPBS and 100µL was pipetted on the coverslips. Coverslips were washed one time with 100µL H_2_O. Coverslips were dried on a paper towel, one drop of mounting media was put on and the coverslips were placed on object plates. The probes were dried overnight at room temperature and then stored at 4°C until microscopy. Pictures were recorded using SP8 confocal microscope (Leica) and analysed with ImageJ.

**Immune phenotyping**

1x10^6^ J774A.1 cells in 1mL media were plated out on a 6 well plate and treated with PPP inhibitors. After 24hrs of incubation, cells were scraped of. Cells were washed with 1mL DPBS. Cells were re-suspended in 90µL DPBS and 10µL murine FcR-blocking agent. Cells were incubated for 10min at 4°C. 900µL DPBS were added and probes were split into 100µL portions. Master mixes of different stains were prepared and added to the cells. Cells were incubated for 20min at 4°C and washed with 1mL DPBS afterwards. For washing, cells were incubated with DPBS for 5min and centrifuged at 300g for 5min afterwards. Cells were re-suspended in 200µL DPBS and measured using Miltenyi MacsQuant X flow cytometer immediately after an initial multi-colour compensation procedure.

For intracellular marker, Fix & Perm Cell Permeabilization kit was used following manufacture´s protocol.

Primary murine macrophages were processed equally.

|  | | | **Macrophage phenotypic marker** | | | | **Isotype control** | | | |
| --- | --- | --- | --- | --- | --- | --- | --- | --- | --- | --- |
| **Laser** | **Channel (nm)** | **Colour** | **1** | **2** | **3** | **4** | **1** | **2** | **3** | **4** |
| 405 | 450/50 | Brilliant violet | CD80 | PD-1 | CCR7 | F4/80 | Hamster IgG | Rat IgG2aκ | Rat IgG2aκ | Rat IgG2aκ |
| 488 | 525/50 | FITC | F4/80 | CD200R | F4/80 | *CD68* | Rec. human IgG1 | Rec. human IgG1 | Rec. human IgG1 | *Rec. human IgG1* |
|  | 585/40 | PE | CD206 | F4/80 | CD86 | *Egr-2* | Rat IgG2aκ | Rat IgG2aκ | Rat IgG2bκ | *Rat IgG2aκ* |
|  | 655-730 | PerCP-Cy5.5 | MHC II | CD64 |  | *iNOS* | Rat IgG2bκ | Mouse IgG1κ |  | *Rat IgG2aκ* |
|  | 750LP | PE-Cy^7^ | CD38 | PD-L1 | CD115 | *IL-10* | Rat IgG2aκ | Rat IgG2b | Rat IgG2aκ | *Rat IgG2bκ* |
| 635 | 655-730 | APC | CD16/32 | TGF-β | SIRPα | *Arg.1* | Rat IgG2aλ | Mouse IgG1κ | Mouse IgG2aκ | *Rat IgG2aκ* |
| 635 | 750LP | APC-Cy^7^ | CD11b | CD11b | CD11b | CD11b | Rat IgG2bκ | Rat IgG2bκ | Rat IgG2bκ | Rat IgG2bκ |

**Immunohistochemical staining of murine spleen**

After preparation of survival cohort of hMB transfected NSG mice after treatment with alemtuzumab and/or S3, a 2x2mm piece of spleen was fixated in formaldehyde. Histological sections were produced and immunohistochemical staining of CD19 and CD68 was performed. Whole slide scans were saved and analysed with NDP.view2 Plus Image viewing software.

***in vivo* experiments**

Survival analysis of hMB transfected NSG mice under treatment with alemtuzumab and/or S3

8-18 week old NSG mice got intravenous injection of 1x10^6^ hMB cells in 100µL PBS.

Four cohorts were built:

1) vehicle + vehicle
2) vehicle + alemtuzumab
3) vehicle + S3
4) alemtuzumab + S3

Three days after tumor cell injection, cohort 3 and 4 were treated for ten days with 20 mg/kg S3 in 200µL 30% PEG 400/0,5% Tween 80/5% propylene glycol (vehicle 200μl 30% PEG 400 /0.5% Tween 80/5% propylene glycol) intraperitoneally. On day 8 after tumor cell injection alemtuzumab was applicated for three days intraperitoneally. On day 8 the mice were injected with alemtuzumab 1mg/kg, on day 9 and 10 with 5mg/kg, in a total volume of 50µL PBS (vehicle 50µL PBS). Mice were scored daily with a score sheet developed for hMB-transfected NSG mice.

Macrophage function analysis of primary murine macrophages of C57BL/6 after treatment with S3

8-18 week old C75BL/6 mice were treated for seven days with 20mg/kg S3 in 200µL 30% PEG 400/0,5% Tween 80/5% propylene glycol (vehicle 200μl 30% PEG 400 /0.5% Tween 80/5% propylene glycol) intraperitoneally. Mice were scored daily and were sacrificed on day eight. Peritoneal macrophages were collected by peritoneal lavage with DMEM medium. Spleen and femurs were dissected. Spleen were mashed through a 30µM filter with DMEM medium and cell suspension was used for further analysis. Femurs got flushed with DMEM medium to collect bone marrow.

**Knockdown cell production**

Plasmid production

Target sequence oligonucleotides were produced by SIGMA-ALDRICH. For oligonucleotide cloning, 1µL of 1µM oligonucleotide, 2,5µL thermopol polymerase buffer, 2,5µL of 5µM primer EcoR1, 2,5µL of 5µM primer Xho1, 0,5µL dNTPs mix, 0,5µL vent polymerase and 14,5µL H_2_O were mixed and polymerase chain reaction was performed. Oligonucleotides were selected by electrophoretic separation using FAE gel with 1,5% agarose and 5µL GELRed Nucleic Acid Gel stain 10000x. After cutting out oligonucleotide bands under UV-light, oligonucleotides were purified with QIAquick PCR purification kit (kit protocol followed). For oligonucleotide digest, 30µL oligonucleotide solution, 1µL primer EcoR1, 1µL primer Xho1 and 8µL NEBuffer 2 were incubated for 2hrs at 37°C. Oligonucleotides were purified again as described above, oligonucleotide concentration was determined with Quick-load 100bp DNA ladder on gel and oligonucleotides were solved in 30µL elution buffer. For ligation 2,92ng oligonucleotide and 97,08ng vector [MLP plasmid, 7893bp] with 1µL 10x T4 DNA Ligase Reaction Buffer and 1µL T4 DNA ligase was used and filled up with H_2_O to 10µL. Probes were incubated for 1hr at room temperature.

Transformation into competent bacteria

Competent E. coli were thawn on ice. 5µL plasmid per 50µL competent cells were added and incubated for 30min on ice. Heat shock for 110sec at 42°C and recovery for 2min on ice was done. Cells were centrifuged at 6000rpm for 1min, supernatant was removed and 450µL fresh LB media (1% tryptone, 0,5% yeast extract, 1% NaCl, filled up with water) was added. Cells were recovered in shaker for 30min at 37°C and 100µL were plated out on LB media layer (1% tryptone, 0,5% yeast extract, 1% NaCl, 1,5% agar filled up with water) with 100µg/mL ampicillin in a 10cm dish.

Plasmid enrichment and purification

When colonies on bacteria plate were visible by eye, 4-5 colonies per plasmid were picked and expanded overnight in 50mL LB media with 100µg/mL ampicillin in shaker at 37°C. For later use a probe of each colony was saved on a new bacteria plate. Plasmids were purified by using I-Blue Midi Plasmid Kit (following kit protocol).

For sequence verification, 10µL of 100ng plasmid with 4µL MSV-5 primer was sent to LGC Genomics GmbH.

Plasmids with correct sequence were picked from back up bacteria plate and expanded and purified again as described above.

Transfection with phoenix retroviral producer line

8x10^6^ HEK293T cells were plated out on a 10cm dish and incubated overnight. Cells were washed and 10mL DMEM media without any supplements was put on the cells. 25µM cholorquine was added and cells were incubated for 10min at 37°C. 18,5µg plasmid, 3µg pMD2.G, 5µg psPax2 and 99µL 2M CaCl_2_ were mixed and filled up with H_2_O to 790µL. Under mixing with bubble formation, 790µL 2x HEPES buffered saline (280mM NaCl, 50mM HEPES, 1,5mM Na_2_HPO_4_, adjusted pH to 7,05) was added and mixture was added immediately dropwise to the HEK293T cells. Cells were incubated for 4hrs, then media was soaked off, cells were washed with 10mL DPBS and 10mL HEK293T cell media was added. 100µM sodium butyrate was added and cells were incubated. After 24hrs and 48hrs supernatant was collected and centrifuged at 1500rpm for 10min. Supernatant was filtered through a 45µm filter and flow through was used for further steps. New media and sodium butyrate was put on the HEK293T cells.

Virus production was verified with fluorescent microscopy and Lenti-X GoStix Plus.

Infection of J774A.1 macrophages

1x10^5^ J774A.1 cells in 1mL media per well were plated out on a 12 well plate and incubated for 24hrs. 1mL of viral media per well was added and cells were spin at 800CF for 2hrs at 32°C. Cells were incubated for 48hrs. Cell media was changed and cell selection with 2µg/mL puromycin was started. Selection efficacy was verified using Miltenyi MacsQuant X flow cytometer. ≥ 95% GFP-positive macrophages were accepted as pure shRNA-transfected cells.

**Metabolomics**

1x10^7^ J774A.1 in 10mL media were plated out on 10cm dishes and incubated for 24hrs. Dishes were treated with PPP inhibitors and incubated for 24hrs. Cells were scraped, washed with DPBS and re-suspended in 1mL DPBS. The cells were counted using CASY cell counter and analyser. Cells were centrifuged at 300g for 5min and supernatant was discarded. Cell pellet was crash frozen using liquid nitrogen. The cell pellets were sent to CEMBIO researcher/ Department of Basic Medical Sciences at San Pablo-CEU University in Madrid (Spain) where the metabolomics analysis was performed in the lab of Dr. Coral Barbas Arribas.

**Maturation staining of primary murine macrophages**

For LSK compartment analysis, ACK lysis buffer was added to bone marrow cells for 2min to lysate erythrocytes and afterwards 5x10^6 cells per mice were hand over for further processing to Felix Picard (AG Holger Winkels, University of Cologne). Additionally, bone marrow cells were processed as described in section Immune phenotyping and stained with maturation antibody panel. Common myeloid progenitor cells: CD41, CD34. Monocytes: CD11b, CX3CR1, Ly6C. Macrophages: CD11b, F4/80, CD64.

**Primary human macrophages**

Isolation of CD14^+^ cells

Peripheral blood mononuclear cells (PBMCs) were isolated from buffy coats of healthy donors provided by blood bank of University Hospital of Cologne. 15mL Lymphoprep was added to a 50mL SepMateTM tube, buffy coats were diluted 1:1 in sterile room temperature DPBS and 20-25mL of it was layered on the top of the Lymphoprep through pipetting it to the walls of the SepMateTM tube. Tubes were centrifuged at 1200g for 15min at room temperature. PBMCs ring were harvested by pouring entire top layer in new 50mL falcon tube. Cells were washed three times with 50mL DPBS and centrifuged at 1300rpm for 8min. Cell pellet was re-suspended in 12mL 4°C MACS buffer (2mM EDTA + 5% Bovine serum albumin (BSA) (PAA Laboratories)), transferred to 15mL tube, centrifuged at 1300rpm for 8min at 4°C and discarded. 200µL of CD14 anti-human magnetically labelled MicroBeads and 800µL of 4°C MACS buffer was added. Magnetic separation was performed by Miltenyi Biotec CD14 human microbead isolation protocol using Miltenyi MacsQuant X flow cytometer. CD14^+^ cells were then re-suspended in RPMI 1640 media. Differentiation was started immediately.

Differentiation of human monocytes under PPP inhibition

1x10^6^ CD14^+^ cells in 2mL RPMI 1640 media per well were plated out on a 12 well plate. 10ng/mL M-CSF and oxythiamine were added at day 1, 3 and 5 without changing the medium. On day 7 the cells were scraped off and used for further experiments.

**Phosphoproteomics**

Sample preparation and lysis

Three dense 25cm dishes with J774A.1 per condition were treated with PPP inhibitors. After 24hrs incubation, plates were washed with 20mL 4°C DPBS per plate. Cell dishes were placed on ice, 1mL RIPA buffer with 1% phosphatase and 1% protease inhibitor was added, cells were scraped off and transferred into 2mL Eppendorf tubes on ice. Cells were centrifuged at full speed for 30min at 4°C. DNA was sheared by sonication with Bioruptor for 10min at 4°C. Samples were centrifuged at full speed for 10min at 4°C and supernatant was transferred into new Eppendorf tubes. Protein concentration was determined by BCA assay. 3mg protein was used for further steps.

Acetone precipitation

Four times volume of cold 100% acetone was added and samples were incubated overnight at -20°C. Samples were centrifuged at 15000 g for 10min at 4°C. Supernatant was discarded and pellet washed twice with 250µL 80-90% acetone under centrifugation at 15000g for 10min at 4°C. Uncapped tubes were left at room temperature for 5-10min to let remaining acetone evaporate without overdrying. Pellet was dissolved in 300µL 6M Urea/2M Thiourea.

In solution digest

1mM 1,4-Dithiothreit (DTT) was added and samples were incubated for 1hr at room temperature. 5,5mM iodoacetic acid (IAA) was added and samples were incubated for 20-45min at room temperature in the dark. 60µL 0,5µg/µL endoprotease Lys-C was added and samples were incubated for 3hrs at room temperature. 900µL 50mM ammonium bicarbonate and 60µL 0,5µg/µL trypsin were added and samples were incubated overnight at room temperature. Samples were acidified with 1% trifluoroacetic acid and centrifuged for 10min at full speed. Supernatant was transferred to a new Eppendorf tube for further steps.

Sample purification by stage tips

C18 columns (200mg Sep Pak of capacity up to 10mg protein) were prepared. Columns were activated with 1mL 100% acetonitrile and washed twice with 1mL 0,1% trifluoroacetic acid. Flow through was discarded and samples loaded on column. Flow through was discarded and column washed twice with 1mL 0,1% trifluoroacetic acid. Peptides were eluted into Eppendorf tubes by adding two times 0,2mL elution buffer (60% acetonitrile, 0,1% formic acid).

Sample enrichment

Phosphopeptide samples were enriched using High select TiO_2_ Phosphopeptide Enrichment Kit by following kit protocol.

Sample measurement

Label-free quantification of peptides was performed on mass spectrometer (Q Exactive^TM^ Plus Hybrid Quadrupole-Orbitatrap^TM^ Mass Spectrometer + EASY-nLC ^TM^ 1200 System) by cooperating CECAD proteomics facility, University of cologne.

Analysis

Raw data acquired from CECAD proteomics facility were filtered and processed on MaxQuant software (v.1.5.3.8) and Perseus software (v.1.5.5.3).

**Proteomics**

Sample preparation and lysis

1x10^7^ J774A.1 cells ins 10mL media per 10cm dish were plated out and after 24hrs treated with PPP inhibitors. After another 24hrs, plates were washed with 10mL 4°C DPBS per plate. Cell dishes were placed on ice, 500µL RIPA buffer with 1% phosphatase and 1% protease inhibitor was added, cells were scraped off and transferred into 2mL Eppendorf tubes on ice. Cells were centrifuged at full speed for 30min at 4°C. DNA was sheared by sonication with Bioruptor for 10min at 4°C. Samples were centrifuged at full speed for 10min at 4°C and supernatant was transferred into new Eppendorf tubes. Protein concentration was determined by BCA assay. 30µg protein was used for further steps.

Acetone precipitation

Four times volume of cold 100% acetone was added and samples were incubated overnight at -20°C. Samples were centrifuged at 15000g for 10min at 4°C. Supernatant was discarded and pellet washed twice with 250µL 80-90% acetone under centrifugation at 15000g for 10min at 4°C. Uncapped tubes were left at room temperature for 5-10min to let remaining acetone evaporate without overdrying. Pellet was dissolved in 60µL 6M urea/2M thiourea.

In solution digest

3µL 1M DTT was added and samples were incubated for 1hr at room temperature. 3µL 550mM IAA was added and samples were incubated for 20-45min at room temperature in the dark. 0,6µL 0,5µg/µL endoprotease Lys-C was added and samples were incubated for 3hrs at room temperature. 180µL 50mM ammonium bicarbonate and 0,6µL 0,5µg/µL trypsin were added and samples were incubated overnight at room temperature. Samples were acidified with 1% trifluoroacetic acid and centrifuged for 10min at full speed. Supernatant was transferred to a new Eppendorf tube for further steps.

Sample purification by stage tips

Stage tips were prepared by stacking 2 layers of SDB-RPS material in a 200µL pipette tip. Stage tips were equilibrated with 20µL 100% methanol and centrifuged at 2600rpm for 2min. 20µL elution buffer (80% acetonitrile + 0,1% trifluoroacetic acid) was added and tips were centrifuged at 2600rpm for 2min. 20µL washing buffer (0,1% trifluoroacetic acid) was added and tips were centrifuged at 2600rpm for 1min. 100µL of the sample was added and tips centrifuged at 2600 rpm for 5min. Tips were washed with 100µL washing buffer and centrifuged at 2600rpm for 3min once and two times with elution buffer. Stage tips were dried with a syringe and stored at -4°C.

Sample measurement

Peptides were eluted with 30µL 1% ammonia in 60% acetonitrile into 96 well plate and dried using SpeedVac concentrator. Label-free quantification of peptides was performed on a mass spectrometer (Q Exactive^TM^ Plus Hybrid Quadrupole-Orbitatrap^TM^ Mass Spectrometer + EASY-nLC ^TM^ 1200 System) by cooperating CECAD proteomics facility, University of cologne.

Analysis

Raw data acquired from CECAD proteomics facility were filtered and processed on MaxQuant software (v.1.5.3.8) and Perseus software (v.1.5.5.3).

**SeaHorse Analysis**

1x10^5^ J774A.1 cells in 100µL media per well were plated out in the XFe96 cell culture microplate. PPP inhibitors were added and cells were incubated for 24hrs. Agilent Seahorse XFe96 Sensor Cartridge was prepared (following Agilent user guide) and cell culture microplate and sensor cartridge preparation for measurement were done (following Agilent protocol). Sensor cartridge injection ports were filled with 20µL of 1µM oligomycin (port A), 22µL of 0,5µM Carbonyl cyanide-*4*-(trifluoromethoxy)phenylhydrazone (FCCP) (port B) and 25µL of 1µM antimycin A plus 100µM rotenone (port C). Four measurement cycles of basal activity of the cells and three cycles of measurement after each injection were performed by Agilent Seahorse XF Analyzer.

**Viability stain**

7AAD staining was performed for hMB cells and Zombie-NIR staining was performed for J774A.1 cells.

For 7AAD staining, 1,5x10^5^ hMB cells were plated out in 100µL media per well of a 96 well plate. At least duplicates were treated with one concentration of the tested inhibitor. As positive control 10% dimethylsulphoxide (DMSO) was used. After 24hrs of incubation cells were transferred into a 96 well U-bottom plate, washed and 7AAD staining was performed. Therefore, 2µL 7AAD stain and 48µL 1x ABB were added per well, cells were incubated for 20min at 4°C and additionally 50µL 1% ABB was added. Readout was performed immediately using Miltenyi MacsQuant X flow cytometer. Inhibitor concentrations with an amount of viable cells ≥ 90% compared to untreated control was accepted as non-toxic concentration.

For Zombie-NIR staining 1,2x10^6^ J774A.1 cells were plated out in 1mL media in a 12 well plate. Duplicates were treated with one concentration of the tested inhibitor. As positive control 10% DMSO was used. After 24hrs of incubation cells were scraped off, transferred into FACS tubes and Zombie-NIR staining was performed. A dilution of Zombie-NIR staining solution 1:100 in DPBS was used. Readout was performed using Miltenyi MacsQuant X flow cytometer. Inhibitor concentrations with an amount of viable cells ≥ 90% compared to untreated control was accepted as non-toxic concentration.

Additionally, Cell titer glo assay was used to measure viability of J774A.1, THP1 and hMB cells. 1x10^4^ J774A.1 cells or THP1 cells respectively 1,5x10^5^ hMB cells were plated out in 100µL media per well of a 96 well plate. Triplets were treated with one concentration of the tested inhibitor. As positive control 10% DMSO was used. After 18hrs cells were washed two times, were transferred to white 96 well plate and Cell titer glo staining was performed. Readout was performed by fluorescence intensity measurement with FLUOStar OPTIMA. Inhibitor concentrations with an ATP amount of the cells ≥ 90% compared to untreated control was accepted as non-toxic concentration.

**Western Blot Analysis**

3x10^6^ J774A.1 respectively 4,5x10^6^ hMB cells were plated out on a six well plate. The cells were treated and incubated for 18hrs. The cells were washed with 1mL DPBS, discarded and stored on ice. 30µL of RIPA buffer (50mM Tris-HCl pH8, 150mM NaCl, 0,1% SDS, 0,5% DOC, 1% NP-40, filled up with ddH2O) with 1x Phosphatase Inhibitor Cocktail 2 and 1x Protease Inhibitor Cocktail were added and probes centrifuged for 30min at full speed at 4 °C. Supernatant was used for the experiments. BCA-Assay was performed to evaluate protein concentration, measured with FLUOStar OPTIMA. 60µg protein per condition was used, volume filled up with RIPA to 5µL and 5µL Urea added. Probes were incubated at 37 °C for 10min. A 10% separating gel (1,85mL Buffer (1,5mM Tris HCL pH8,8, 0,4% SDS, ddH_2_O), 1,66mL 30% Rotiphorese, 1,5mL ddH_2_O, 40,6µL APS and 4,06µL Temed) and a 5% stacking gel (0,31mL Buffer (0,5M Tris HCl pH6,8, 0,4% SDS, ddH_2_O), 0,42mL 30% Rotiphorese, 1,75mL ddH_2_O, 12,5µL APS and 1,25µL Temed) were produced. 3,5µL Page Ruler Prestained NIR Protein Ladder was used. Western Blot run was performed in Running Buffer (25mM Tris, 192mM glycine, 3,5mM SDS in ddH_2_O) at constant 80V until stacking gel was passed and at constant 150V in separating gel. Gel was blotted on nitrocellulose membrane Hybind-C at constant 400mA for 1hr in transfer Buffer (25mM Tris, 192mM glycine, ddH_2_O). Total Protein stain was performed with REVERT Total protein stain and membrane was blocked with 10mL TBS-T (10mM Tris, 250mM NaCl, HCl pH7,6, 0,05% Tween20, ddH_2_O) + 5% BSA for 1hr. First antibody was diluted in 5mL TBS-T + 5% BSA and membrane was incubated overnight at 4°C in the dark. Membrane was washed to times with 2mL TBS-T for 10min and one time with 2mL TBS (10mM Tris, 250mM NaCl, HCl pH7,6, ddH_2_O) for 10min. Second antibody was diluted in 2,5mL TBS + 2,5mL Odyssey Blocking Buffer. Membrane was incubated with second antibody for 1hr at room temperature. Membrane was washed three times with TBS for 10min. Membrane fluorescence was measured with ODYSSEY CLx. Fluorescence intensity was calculated with Image Studio Lite Vers. 5.2.

**QUANTIFICATION AND STATISTICAL ANALYSIS**

**Statistical analysis**

Statistical analysis was performed using GraphPad Prism software. Significance was calculated using unpaired t-test (Fig. 1multiple comparison one-way ANOVA, two-way ANOVA (Fig. 3F), RM one-way ANOVA (Fig. 6H), paired t-text (Fig. 6 I-L), unpaired t-test (Fig. 7A-E), Benjamini-Hochberg test (Fig. 7F).

In figure 1, figure 2, figure 4G, figure 5B-C and E-F, figure 6A-H, figure 7A-C and E data are shown as mean ± SEM. In figure 3 surface marker stain is shown as mean of four replicates, SeaHorse analysis over time is shown as one representative example mean ± SD, calculated parameters of SeaHorse analysis are shown as mean ± 5-95 percentile. In figure 5D metabolite amount is shown as Min. to Max. In figure 6 I-L data are shown as single data plots. In figure 7C surface marker stain is shown as mean of ten replicates.

Statistical values, including number of replicates (n), are named in the figure legends. In vitro experiments: n = number of separate experiments; *in vivo* experiments: n = number of individual animals. **p* < 0.05; ***p* < 0.01; ****p* < 0.001; *****p* < 0.0001.

**Proteomic and phosphoproteomic analysis**

Raw data acquired from CECAD proteomics facility were filtered and processed on MaxQuant software (v.1.5.3.8) and Perseus software (v.1.5.5.3). Data are generated out of one experiment with three replicates per condition.

Volcano plots of proteomics were generated with software of Bioconductor. The mean of the different inhibitor treatment respectively PPP enzyme knockdowns was calculated and compared to the untreated control. Significance was defined as mean Log_2_ fold change > 0,5 or < -0,5 and q-value <0,05. Circle size represents the number of significant occurrence in the different treatment conditions.

For circle plot analysis, significant genes were extracted from proteomic- and phosphoproteomic analysis. Significance was defined as mean Log_2_ fold change > 0,5 or < -0,5 and q-value <0,05 for proteomic analysis and as mean Log_2_ fold change > 0,5 or < -0,5 and p-value >1,3 for phospho-proteomic analysis. Significant genes were clustered with String analysis. The ten biggest clusters were used for further analysis. Clusters were named by using GeneAnalytics and similar clusters were merged in one heading. Mean of -log_10_ p-value of the clusters was calculated and is represented in heat colour, count of genes per cluster is represented in circle size.

Normalized upstream kinase score out of phosphoproteomic analysis, was calculated by adapted code of INKA analysis. On basis of the work of Beekhof *et al.* ^24^, we calculated a simplified upstream kinase score analysis, using the murine data available from the PhosphoSitePlus (PSP) database:

$Upstream Kinase Score=\surd$∑_Kin_ x ∑_PSP_

With ∑_Kin_ representing the sum of all phosphopeptides observed in the experiment per kinase found in the murine PSP database, whilst ∑_PSP_ representing the sum of all substrate phosphopeptides observed in the experiment associated with each kinase found in the murine PSP database. These scores were calculated for each replicate per condition, with the normalized upstream kinase score (NUKS) representing the mean difference in upstream kinase scores between untreated control and treatment with PPP inhibitors and macrophage wildtype and PPP knockdown macrophages respectively.

**KEY RESOURCE TABLE**

| REAGENT or RESOURCE | SOURCE | IDENTIFIER |
| --- | --- | --- |
| Antibodies | | |
| Rabbit polyclonal anti-ß-actin | BioLegend | Cat# 622101, RRID:AB_315945 |
| Sheep polyclonal anti-Arg1 | RnD Systems | Cat# IC5868P |
| Mouse monoclonal anti-CCR7 (CD197) | BioLegend | Cat# 353213, RRID:AB_10915474 |
| Rat monoclonal anti-CD11b | BioLegend | Cat# 101226, RRID:AB_830642 |
| Rat monoclonal anti-CD115 (CSF-1R) | BioLegend | Cat# 135523, RRID:AB_2566459 |
| Rat monoclonal anti-CD16/32 | BioLegend | Cat# 156607, RRID:AB_2800705 |
| Rat monoclonal anti-CD19 | Thermo Fisher Scientific | Cat# 14-0194-80, RRID:AB_2637170 |
| Mouse monoclonal anti-CD200R | BioLegend | Cat# 329305, RRID:AB_2074201 |
| Mouse monoclonal anti-CD206 | BD Biosciences | Cat# 551135, RRID:AB_394065 |
| Rat monoclonal anti-CD38 | BioLegend | Cat# 102717, RRID:AB_2072892 |
| Mouse monoclonal anti- CD64 | BioLegend | Cat# 305025, RRID:AB_2561587 |
| Rabbit polyclonal anti-CD68 | Abcam | Cat# ab125212, RRID:AB_10975465 |
| Rat monoclonal anti-CD68 | BioLegend | Cat# 137017, RRID:AB_2562949 |
| Mouse monoclonal anti-CD80 | BD Biosciences | Cat# 557227, RRID:AB_396606 |
| Rat monoclonal anti-CD86 | Miltenyi Biotec | Cat# 130-123-724, RRID:AB_2889634 |
| Mouse monoclonal anti-CX3CR1 | BioLegend | Cat#149007,  RRID:AB_2564491 |
| Rat monoclonal anti-EGR-2 | Thermo Fisher Scientific | Cat# 17-6691-82, RRID:AB_11151502 |
| Rat monoclonal anti-F4/80 | BioLegend | Cat# 123110, RRID:AB_893486 |
| Rat monoclonal anti-F4/80 | BioLegend | Cat# 123124, RRID:AB_893475 |
| Human cell line monoclonal anti-F4/80 | Miltenyi Biotec | Cat # 130-102-327, RRID:AB_2651701 |
| Mouse monoclonal anti-human HLA-DR (MHC II) | BioLegend | Cat# 307604, RRID:AB_314682 |
| Mouse [Histofine Simple Stain Mouse MAX-PO](https://scicrunch.org/resources/data/record/nif-0000-07730-1/AB_2819094/resolver?q=%2A&l=&filter%5b%5d=Vendor:nichirei&i=2754550) | Nichirei Biosciences | Cat# 414341F, RRID:AB_2819094 |
| Rat monoclonal anti-IL-10 | BioLegend | Cat# 505025, RRID:AB_11149682 |
| Mouse monoclonal anti-iNOS | Novus Biologicals | Cat# NBP2-22119 |
| Rabbit monoclonal anti-IRF1 (D5E4) | Cell Signaling Technology | Cat# 8478, RRID:AB_10949108 |
| Rabbit polyclonal anti-IRG1 | Cell Signaling Technology | Cat# 17805 |
| Rat monoclonal anti-Ly6c | BioLegend | Cat# 128011, RRID:AB_1659242 |
| Rabbit polyclonal anti-P2RY14 | LSBio | Cat# LS‑C409714-20 |
| Rabbit monoclonal anti-PD-1 (D7D5W) | Cell Signaling Technology | Cat# 84651, RRID:AB_2800041 |
| Rabbit polyclonal anti-PD-L1 | Thermo Fisher Scientific | Cat# PA5-20343, RRID:AB_11153819 |
| Mouse monoclonal anti-phosphogluconate dehydrogenase (G-2) (6PGD) | Santa Cruz Biotechnology, INC. | Cat# sc-398977, RRID:AB_2827766 |
| Rabbit monoclonal anti- Protein-tyrosine kinase 2-beta  (PYK2) | Abcam | Cat# ab32571, RRID:AB_777566 |
| Rabbit polyclonal anti-SIRP1a | SIGMA-ALDRICH | Cat# SAB2102154, RRID:AB_10605073 |
| Rabbit monoclonal anti-STAT1 | Cell Signaling Technology | Cat# 80916, RRID:AB_2799965 |
| Mouse monoclonal anti-TGF-beta1 | BioLegend | Cat# 349706, RRID:AB_10680787 |
| Rabbit polyclonal anti-transketolase (TKT) | Biorbyt Ltd. | Cat# orb247362 |
| Rabbit polyclonal anti-UGP2 | Thermo Fisher Scientific | Cat# PA5-27760, RRID:AB_2545236 |
| Bacterial and virus strains | | |
| 5-alpha Competent E. coli | New England Biolabs Inc. | Cat# C2987H |
| Chemicals, peptides, and recombinant proteins | | |
| 1-hydroxy-8-methoxy-anthraquinone (S3) | SIGMA-ALDRICH | Cat# R164046 |
| 1-hydroxy-8-methoxy-anthraquinone (S3) | SAGECHEM LIMITED | Cat# S474625 |
| 1,4-Dithiothreit | CARL ROTH | Cat# 6908.2 |
| 2-deoxy-D-glucose | SIGMA-ALDRICH | Cat# D8357 |
| 6-aminonicotinamide | SIGMA-ALDRICH | A0630; CAS: 329-89-5 |
| 6-phosphogluconolactone | SIGMA-ALDRICH | Cat# P7877 |
| Alemtuzumab | MabCampath | NDC code 58468-0357-3 |
| Bendamustine hydrochloride hydrate | SIGMA-ALDRICH | Cat# B5437 |
| BML-275 hydrochloride | SIGMA-ALDRICH | Cat# **ADVH7F38323F** |
| DAPI (4′,6-Diamidino-2-phenylindoldihydrochlorid) | SIGMA-ALDRICH | Cat# D9542 |
| Daratumumab | Janssen-Cilag International N.V. | EMEA/H/C/004077 |
| D-erythrose-4-phosphate sodium | SIGMA-ALDRICH | E0377; CAS: 103302-15-4 |
| D-fructose-6-phosphate disodium salt hydrate | SIGMA-ALDRICH | F3627; CAS: 26177-86-6 |
| D-glucose-6-phosphate sodium salt | SIGMA-ALDRICH | Cat# G7879 |
| dNTP mix | Thermo Fisher Scientific | Cat# R0192 |
| D-ribose-5-phosphate disodium salt hydrate | SIGMA-ALDRICH | R7750; CAS: 18265-46-8 |
| D-ribulose-5-phosphate sodium salt | SIGMA-ALDRICH | Cat# **R9875** |
| D-xylulose-5-phosphate lithium salt | SIGMA-ALDRICH | Cat# 78963 |
| DL-glyceraldehyde-3-phosphate | SIGMA-ALDRICH | G5251; CAS: 591-59-3 |
| LentiX GoStix Plus | Takara Bio Inc. | Cat# 631280 |
| Lymphoprep | STEMCELL Technologies | Cat# 07801 |
| M-CSF, recombinant human | Thermo Fisher Scientific | Cat# **PHC9501** |
| M-CSF, recombinant mouse | Thermo Fisher Scientific | Cat# **PMC2044** |
| Microbeads CD14 human | Miltenyi Biotec | Cat# 130-050-201, RRID:AB_2665482 |
| Obinutuzumab | Ro­che Re­gis­tra­ti­on Li­mi­ted | EMEA/H/C/002799 |
| Oligomycin | SIGMA-ALDRICH | Cat# **O4876** |
| Oxythiamine chloride hydrochloride | SIGMA-ALDRICH | Cat# O4000 |
| PageRuler prestained NIR Protein ladder | Thermo Fisher Scientific | Cat# 26635 |
| P-hydroxyphenylpyruvate 98% | SIGMA-ALDRICH | 114286; CAS: 156-39-8 |
| Physcion | SIGMA-ALDRICH | 17797; CAS: 521-61-9 |
| Restriction Endonuclease EcoR1 | New England Biolabs Inc. | Cat# R0101L |
| Restriction Endonuclease Xho1 | New England Biolabs Inc. | Cat# R0146L |
| Vent polymerase | New England Biolabs Inc. | Cat# M0245S |
| Critical commercial assays | | |
| 7AAD viability staining eBioscience | Thermo Fisher Scientific | Cat# A1310 |
| BCA Protein Assay Kit | Thermo Fisher Scientific | Cat# 23227 |
| CellTiter-Glo Luminescent Cell Viability Assay Kit | Promega | Cat# G7570 |
| Fix & Perm Cell Permeabilization Kit | Thermo Fisher Scientific | Cat# GAS003 |
| Human IL-6 ELISA MAX Standard Set |  | Cat# 430501 |
| Human IL-10 ELISA MAX Standard Set |  | Cat# 430601 |
| I-Blue Midi Plasmid Kit | IBI SCIENTIFIC | Cat# IB47180 |
| Mouse IL-6 ELISA MAX Standard Set | BioLegend | Cat# 431301, RRID:AB_2883997 |
| Mouse IL-10 ELISA MAX Standard Set | BioLegend | Cat# 431411 |
| Odyssey Blocking Buffer | LI-COR | Cat# 927-40000 |
| Phosphopeptide Enrichment Kit | Thermo Fisher Scientific | Cat# A32993 |
| QIA quick PCR Purification Kit | QIAGEN | Cat# 28104 |
| REVERT Total protein stain | LI-COR Biotech. | Cat# 926-11011 |
| SeaHorse XF Base Medium | Agilent Technologies, Inc. | Cat# 103334-100 |
| SeaHorse XFe96 FluxPak | Agilent Technologies, Inc. | Cat# 102416-100 |
| Zombie NIR Fixable Viability Kit | BioLegend | Cat# 423105 |
| Deposited data | | |
| Affinity-based mass spectrometry performed with 5680 proteins (Proteomic analysis) | This paper | Accession number PXD042428, https://www.ebi.ac.uk/pride/ |
| Affinity-based mass spectrometry performed with 19383 protein-sites (Phospho-proteomic analysis) | This paper | Accession number PXD042428, https://www.ebi.ac.uk/pride/ |
| Murine database for phosphopeptides | PhosphoSitePlus | https://www.phosphosite.org/staticDownloads |
| Experimental models: Cell lines | | |
| Human: HEK293T-CAF40-null | DSMZ | Cat# ACC-872, RRID:CVCL_A5EE |
| Human: THP-1 | DSMZ | Cat# ACC-16, RRID:CVCL_0006 |
| Humanized mouse cells: hMB, strain 102 |  | Leskov, I. *et al.* Rapid generation of human B-cell lymphomas via combined expression of Myc and Bcl2 and their use as a preclinical model for biological therapies. *Oncogene* 32, 1066–1072 (2013). |
| Mouse: J774A.1 | ATCC | Cat# TIB-67,  RRID:CVCL_0358 |
| Human: L-929 | DSMZ | Cat# ACC-2,  RRID:CVCL_0462 |
| Experimental models: Organisms/strains | | |
| Mouse: C57BL/6J | Jackson laboratory | Cat# 000664, RRID:IMSR_JAX:000664 |
| Mouse: Wild-type NOD.Cg-Prkdc^scid^ Il2rg^tm1Wjl^/SzJ (NSG) | Jackson laboratory | Cat# 005557/NSG, RRID:IMSR_JAX:005557 |
| Oligonucleotides | | |
| 6PGD_1 | SIGMA-ALDRICH | Oligo# 8810932277-000060 |
| 6PGD_2 | SIGMA-ALDRICH | Oligo# 8810932277-000070 |
| TKT_1 | SIGMA-ALDRICH | Oligo# 8810932277-000040 |
| TKT_2 | SIGMA-ALDRICH | Oligo# 8810932277-000050 |
| Software and algorithms | | |
| Enhanced Volcanoplot software (Figure 4) | Bioconductor | https://bioconductor.org/packages/release/bioc/html/EnhancedVolcano.html |
| FlowJo 10.7.1 | FlowJo | https://www.flowjo.com/solutions/flowjo |
| GeneAnalytics | LifeMap Sciences | https://geneanalytics.genecards.org/ |
| GraphPad Prism6 | GraphPad Software | https://www.graphpad.com/ |
| Image Studio Lite | LI-COR Biotechnology | https://www.licor.com |
| ImageJ | [U.S. Department of Health & Human Services](https://www.hhs.gov/) | https://imagej.nih.gov/ij/download.html |
| INKA (original and own mouse modification) | Molecular System Biology; this paper | Beekhof, R. *et al.* INKA , an integrative data analysis pipeline for phosphoproteomic inference of active kinases . *Mol. Syst. Biol.* 15, (2019).; |
| MACSQuantify | Miltenyi Biotec | https://www.miltenyibiotec.com |
| MaxQuant | Max-Planck-Institute of Biochemistry | https://maxquant.net/maxquant/ |
| NDP.view2 Plus Image viewing software U12388-02 | Hamamatsu Photonics Deutschland GmbH | https://www.hamamatsu.com/eu/en/product/life-science-and-medical-systems/digital-slide-scanner/U12388-01.html |
| Perseus | Max-Planck-Institute of Biochemistry | https://maxquant.net/perseus/ |
| PRIDE database | EMBL-EBI | https://www.ebi.ac.uk/pride/ |
| StringAnalysis | STRING Consortium 2021 | https://string-db.org |
