## Supplements for "Pentose Phosphate Pathway Inhibition activates Macrophages towards phagocytic Lymphoma Cell Clearance"

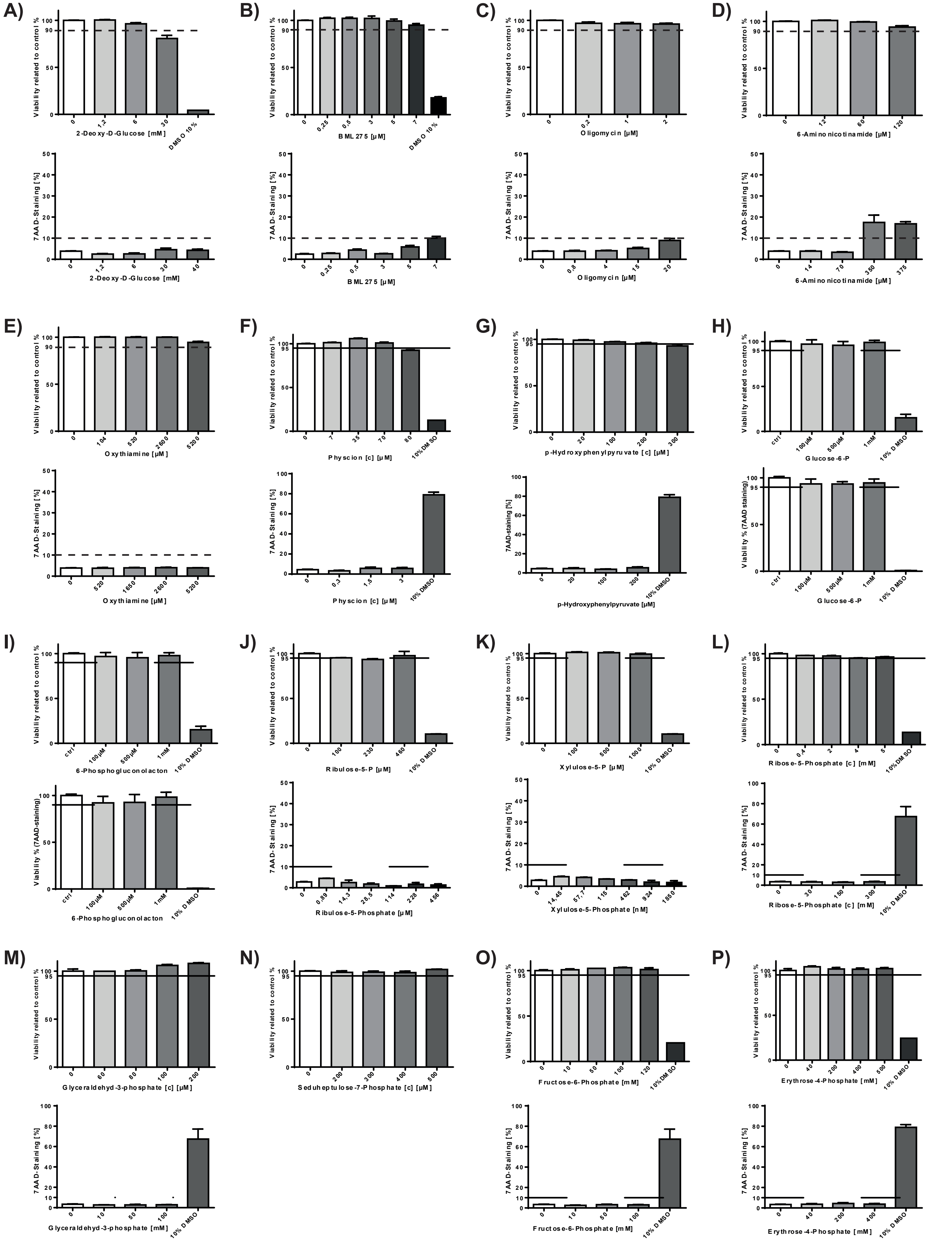
A)
Viability related to control %
2-Deoxy-D-Glucose [mM]
DMSO 10%
7AAD-Staining [%]
2-Deoxy-D-Glucose [mM]
B)
BML 275 [µM]
DMSO 10%
BML 275 [µM]
C)
Oligomycin [µM]
Oligomycin [µM]
D)
6-Aminonicotinamide [µM]
6-Aminonicotinamide [µM]
E)
Oxythiamine [µM]
Oxythiamine [µM]
F)
Physcion [c] [µM]
10% DMSO
Physcion [c] [µM]
G)
p-Hydroxyphenylpyruvate [c] [µM]
7AAD-staining [%]
p-Hydroxyphenylpyruvate [µM]
H)
Glucose-6-P
Viability % (7AAD staining)
Glucose-6-P
I)
6-Phosphogluconolacton
Viability % (7AAD-staining)
6-Phosphogluconolacton
J)
Ribulose-5-P [µM]
Ribulose-5-Phosphate [µM]
K)
Xylulose-5-P [µM]
Xylulose-5-Phosphate [nM]
L)
Ribose-5-Phosphate [c] [mM]
Ribose-5-Phosphate [c] [mM]
M)
Glyceraldehyd-3-phosphate [c] [µM]
Glyceraldehyd-3-phosphate [mM]
N)
Seduheptulose-7-Phosphate [c] [µM]
O)
Fructose-6-Phosphate [mM]
Fructose-6-Phosphate [mM]
P)
Erythrose-4-Phosphate [mM]
Erythrose-4-Phosphate [mM]

**Suppl. Fig.1 Evaluation of cytotoxicity of used compounds in J774A.1 macrophages and hMB cells.**
**(A-P)** Measurement of viable cells under treatment with different inhibitors. Treatment with 10% DMSO used as positive control. Viability of J774A.1 cells (upper plots) was determined by Zombie staining, viability of hMB cells was determined by 7AAD staining. Viability under inhibition was compared to viability of untreated control cells. Used Inhibitors **A** 2-deoxy-D-glucose, **B** BML275, **C** oligomycin, **D** 6-aminonicotinamide, **E** oxythiamine, **F** physcion, **G** p-hydroxyphenylpyruvate, **H** glucose-6-phosphate, **I** 6-phosphogluconolactone, **J** ribulose-6-phosphate, **K** xylulose-5-phosphate, **L** ribose-5-phosphate, **M** glyceraldehyde-3-phosphate, **N** sedoheptulose-7-phosphate, **O** fructose-6-phosphate, **P** erythrose-4-phosphate.
n=2-6. Data are shown as mean ± SEM. P values were calculated using one-way ANOVA.
*$p < 0.05$; **$p < 0.01$; ***$p < 0.001$; ****$p < 0.0001$.

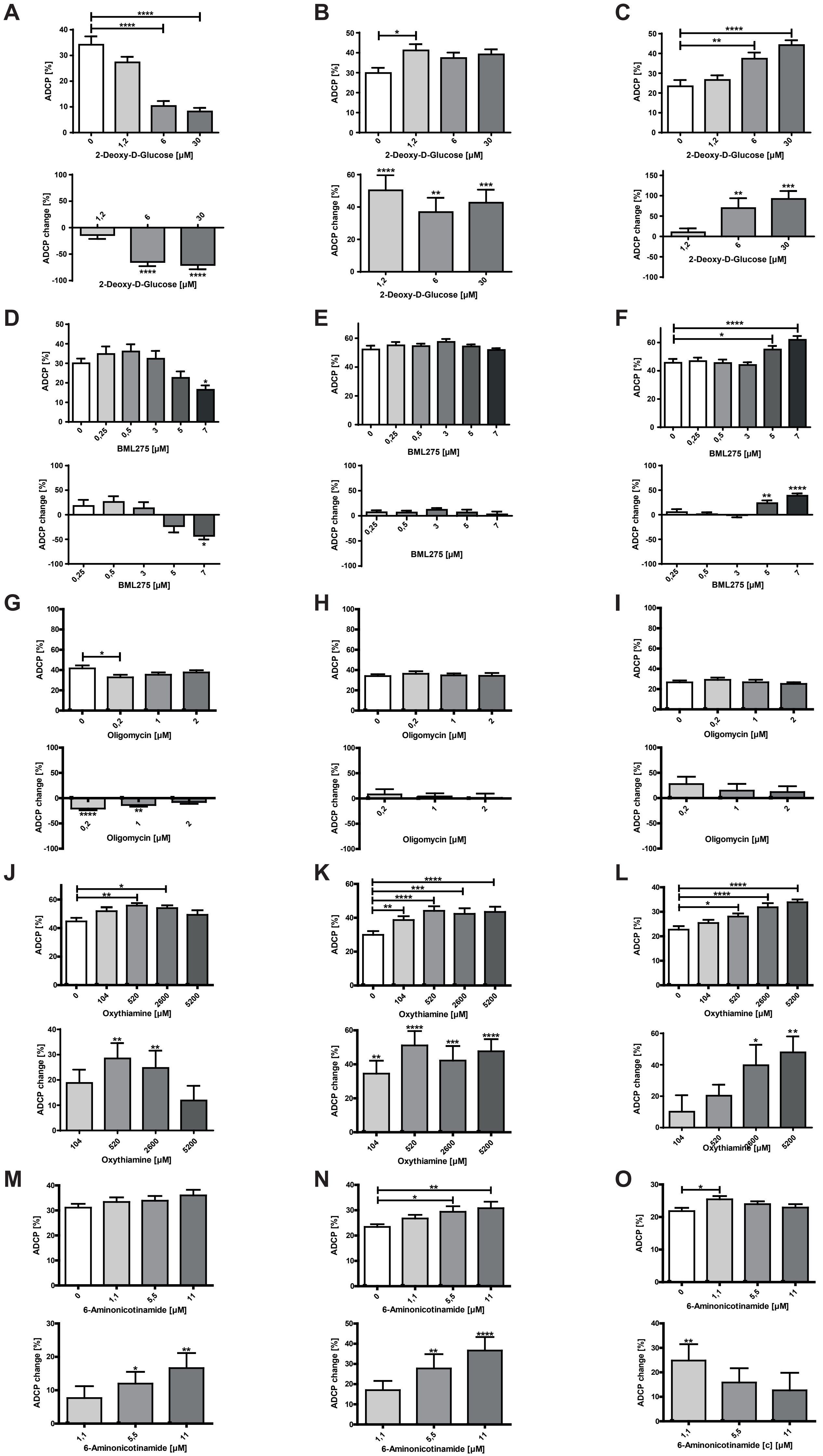

A
ADCP [%]
0
1,2
6
30
2-Deoxy-D-Glucose [µM]
****
ADCP change [%]
B
ADCP [%]
2-Deoxy-D-Glucose [µM]
ADCP change [%]
2-Deoxy-D-Glucose [µM]
C
ADCP [%]
2-Deoxy-D-Glucose [µM]
ADCP change [%]
2-Deoxy-D-Glucose [µM]
D
ADCP [%]
0
0,25
0,5
3
5
7
BML275 [µM]
ADCP change [%]
BML275 [µM]
E
ADCP [%]
BML275 [µM]
ADCP change [%]
BML275 [µM]
F
ADCP [%]
BML275 [µM]
ADCP change [%]
BML275 [µM]
G
ADCP [%]
0
0,2
1
2
Oligomycin [µM]
ADCP change [%]
Oligomycin [µM]
H
ADCP [%]
Oligomycin [µM]
ADCP change [%]
Oligomycin [µM]
I
ADCP [%]
Oligomycin [µM]
ADCP change [%]
Oligomycin [µM]
J
ADCP [%]
0
104
520
2600
5200
Oxythiamine [µM]
ADCP change [%]
Oxythiamine [µM]
K
ADCP [%]
Oxythiamine [µM]
ADCP change [%]
Oxythiamine [µM]
L
ADCP [%]
Oxythiamine [µM]
ADCP change [%]
Oxythiamine [µM]
M
ADCP [%]
0
1,1
5,5
11
6-Aminonicotinamide [µM]
ADCP change [%]
6-Aminonicotinamide [µM]
N
ADCP [%]
6-Aminonicotinamide [µM]
ADCP change [%]
6-Aminonicotinamide [µM]
O
ADCP [%]
6-Aminonicotinamide [µM]
ADCP change [%]
6-Aminonicotinamide [c] [µM]

**Suppl. Fig.2 Metabolic modulation changes antibody-dependent cellular phagocytosis (ADCP) of hMB cells by macrophages.**
**(A-O)** ADCP rate and ADCP rate compared to basal phagocytosis rate (=ADCP change) under treatment with metabolic inhibitors in a co-culture of J774A.1 macrophages and hMB cells under antibody treatment with alemtuzumab. **A**, **D**, **G**, **J**, **M** J774A.1 macrophages pre-treated with metabolic inhibitor, **B**, **E**, **H**, **K**, **N** inhibitor treatment of the co-culture, **C**, **F**, **I**, **L**, **O** hMB cells pre-treated with metabolic inhibitor. Used inhibitors **A-C** 2-deoxy-D-glucose, **D-F** BML275, **G-I** oligomycin, **J-L** oxythiamine, **M-O** 6-aminonicotinamide.
n=3-6. Data are shown as mean ± SEM. P values were calculated using one-way ANOVA.
*p < 0.05; **p < 0.01; ***p < 0.001; ****p < 0.0001.

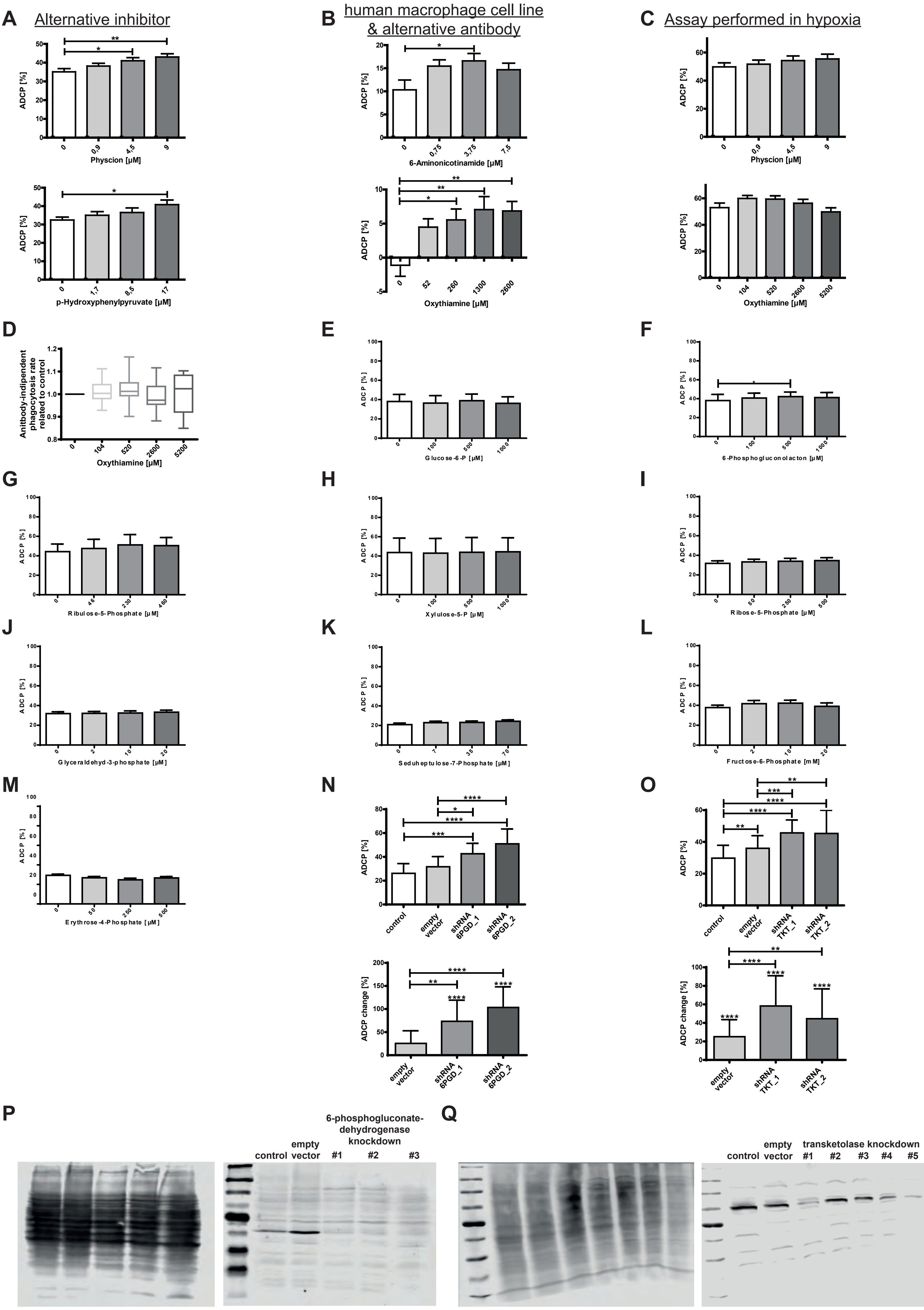
A
Alternative inhibitor
ADCP [%]
Physcion [µM]
p-Hydroxyphenylpyruvate [µM]
B
human macrophage cell line & alternative antibody
6-Aminonicotinamide [µM]
Oxythiamine [µM]
C
Assay performed in hypoxia
Physcion [µM]
Oxythiamine [µM]
D
Anitbody-indipendent phagocytosis rate related to control
Oxythiamine [µM]
E
Glucose-6-P [µM]
F
6-Phosphogluconolacton [µM]
G
Ribulose-5-Phosphate [µM]
H
Xylulose-5-P [µM]
I
Ribose-5-Phosphate [µM]
J
Glyceraldehyd-3-phosphate [µM]
K
Seduheptulose-7-Phosphate [µM]
L
Fructose-6-Phosphate [mM]
M
Erythrose-4-Phosphate [µM]
N
control
empty vector
shRNA 6PGD_1
shRNA 6PGD_2
ADCP change [%]
O
shRNA TKT_1
shRNA TKT_2
P
6-phosphogluconate-dehydrogenase knockdown
control
empty vector
#1
#2
#3
Q
transketolase knockdown
#4
#5

**Suppl. Fig.3 PPP modulation changes ADCP of hMB cells by macrophages.**
**(A, C)** ADCP rate and ADCP rate compared to basal phagocytosis rate (=ADCP change) under treatment with metabolic inhibitors in a co-culture of J774A.1 macrophages and hMB cells under antibody treatment with alemtuzumab. **A** ADCP performed under PPP inhibition with physcion or p-hydroxyphenylpyruvate, **B** ADCP performed with THP1 monocytes and hMB cells under antibody treatment with obinutuzumab and PPP inhibition with 6-aminonicotinamide or oxythiamine. **C** ADCP assay performed in hypoxia under PPP inhibition with physcion or oxythiamine. **D** antibody-independent cellular phagocytosis (AICP) rate of hMB cells by J744A.1 macrophages under treatment with oxythiamine. **(E-M)** ADCP rate under supplementation of PPP intermediates.
**E** glucose-6-phosphate, **F** 6-phosphogluconolactone, **G** ribulose-5-phosphate, **H** xylulose-5-phosphate, **I** ribose-5-phosphate, **J** glyceraldehyde-3-phosphate, **K** sedoheptulose-7-phosphate, **L** fructose-6-phosphate, **M** erythrose-4-phosphate. **(N-O)** ADCP rate and ADCP rate compared to basal phagocytosis rate (=ADCP change) of hMB cells by shRNA mediates PPP knockdown macrophages. **N** shRNA mediated knockdown of 6-phosphogluconate dehydrogenase, **O** shRNA mediated knockdown of transketolase. **(P)** One representative example of western blot analysis of J744A.1 macrophages transfected with empty vector control and shRNA targeting 6-phosphogluconate dehydrogenase. Total protein stain and staining of 6-phosphogluconate dehydrogenase. **(Q)** One representative example of western blot analysis of J744A.1 macrophages transfected with empty vector control and shRNA targeting transketolase. Total protein stain and staining of transketolase.
n=3-6. In **A-O** data are shown as mean ± SEM. P values were calculated using one-way ANOVA.
*$p < 0.05$; **$p < 0.01$; ***$p < 0.001$; ****$p < 0.0001$.

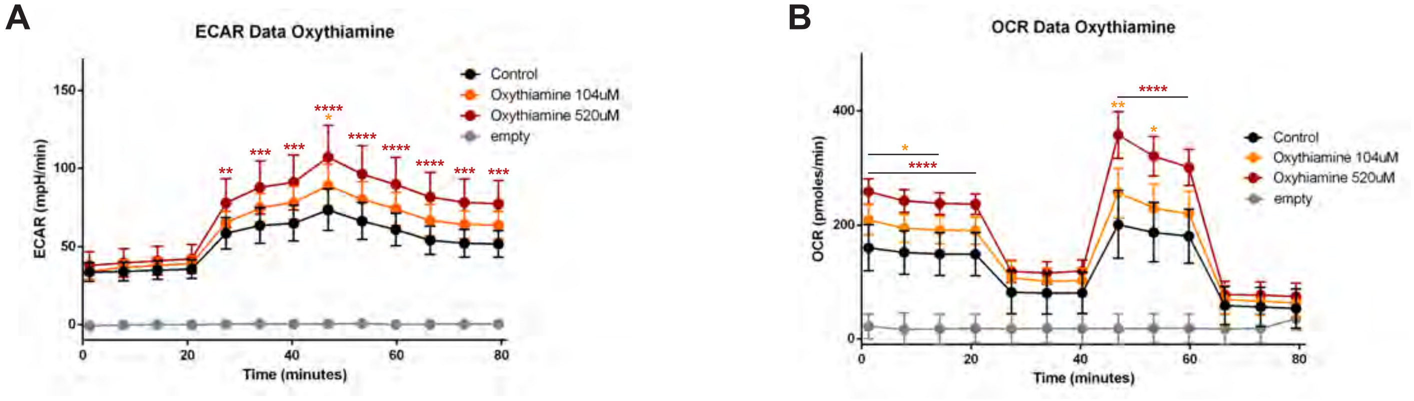


**Suppl. Fig.4 PPP modulation increases metabolic activity in macrophages.**

**(A-B)** Measurement of metabolic activity of J774A.1 macrophages under drug mediated PPP inhibition with oxythiamine by SeaHorse analysis. **A** one representative example of MitoStress test measurement of ECAR, **B** one representative example of MitoStress test measurement of OCR.

Data are shown as mean of six replicates in one experiment ± SD, n=6. P values were calculated using one-way ANOVA.

*p < 0.05; **p < 0.01; ***p < 0.001; ****p < 0.0001.

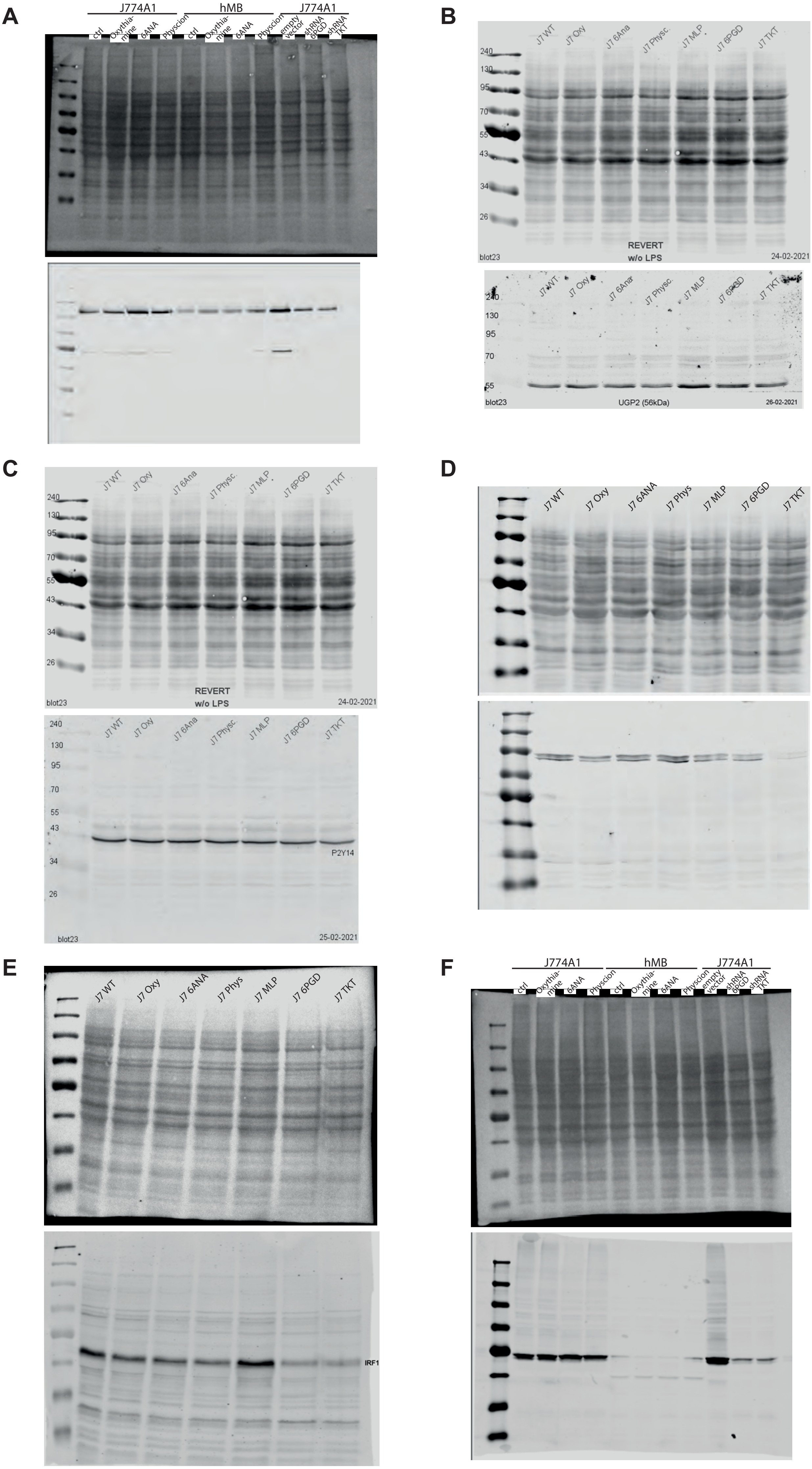
A
J774A1
hMB
J774A1
ctrl
Oxythia-mine
6ANA
Physcion
ctrl
Oxythia-mine
6ANA
Physcion
empty vector
shRNA 6PGD
shRNA TKT
B
J7 WT
J7 Oxy
J7 6Ana
J7 Physc.
J7 MLP
J7 6PGD
J7 TKT
240
130
95
70
55
43
34
26
blot23
REVERT
w/o LPS
24-02-2021
J7 WT
J7 Oxy
J7 6Ana
J7 Physc.
J7 MLP
J7 6PGD
J7 TKT
240
130
95
70
55
blot23
UGP2 (56kDa)
26-02-2021
C
J7 WT
J7 Oxy
J7 6Ana
J7 Physc.
J7 MLP
J7 6PGD
J7 TKT
240
130
95
70
55
43
34
26
blot23
REVERT
w/o LPS
24-02-2021
240
130
95
70
55
43
34
26
J7 WT
J7 Oxy
J7 6Ana
J7 Physc.
J7 MLP
J7 6PGD
J7 TKT
P2Y14
blot23
25-02-2021
D
J7 WT
J7 Oxy
J7 6ANA
J7 Phys
J7 MLP
J7 6PGD
J7 TKT
E
J7 WT
J7 Oxy
J7 6ANA
J7 Phys
J7 MLP
J7 6PGD
J7 TKT
IRF1
F
J774A1
hMB
J774A1
ctrl
Oxythia-mine
6ANA
Physcion
ctrl
Oxythia-mine
6ANA
Physcion
empty vector
shRNA 6PGD
shRNA TKT

**Suppl. Fig.5 PPP inhibition changes the protein expression of hypothesized metabolic-immune response axis in macrophages.**

**(A-F)** One representative example of western blot analysis of J744A.1 macrophages after drug mediated inhibition of the PPP or shRNA mediated knockdown of the PPP. Total protein stain and staining of protein of interest are shown. In **A** and **F** also hMB cells under PPP inhibition has been tested. **A** PYK2 staining, **B** UGP2 staining, **C** P2Y14 staining, **D** STAT1 staining, **E** IRF1 staining, **F** IRG1 staining.

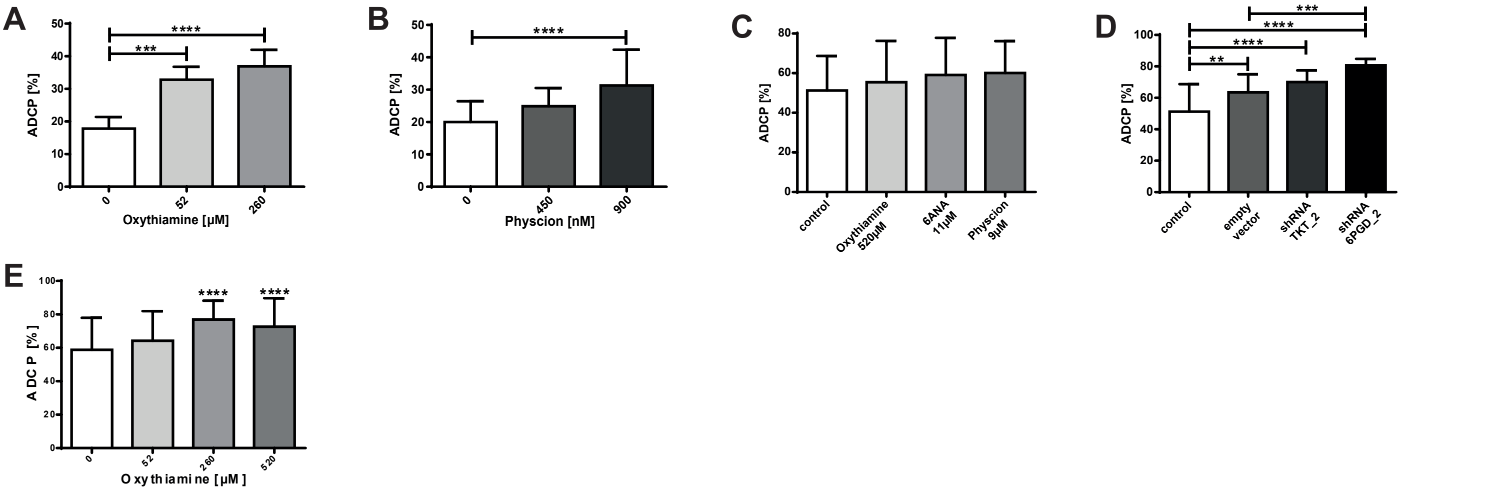


**Suppl. Fig.6 PPP inhibition in primary human environment increases phagocytic capacity of macrophages.**
**(A-B)** ADCP rate of primary human monocyte derived macrophages differentiated in the presence of PPP inhibitors. **A** ADCP change by primary monocyte derived macrophages differentiated in the presence of physcion and M-CSF. **B** ADCP change by primary monocyte derived macrophages differentiated in the presence of oxythiamine and M-CSF. **(C-D)** ADCP rate of primary CLL patient cells by J774A.1 macrophages. **C** ADCP rate under drug mediated PPP inhibition, **D** ADCP rate under shRNA mediated PPP knockdown. **(E)** ADCP rate of primary CLL patient cells by primary human monocyte derived macrophages differentiated in the presence of oxythiamine and M-CSF.
**A** n=5, **B** n=6, **C-D** n=4, **E** n=13. Data are shown as mean ± SEM. P values were calculated by using one-way ANOVA.
*$p < 0.05$; **$p < 0.01$; ***$p < 0.001$; ****$p < 0.0001$.

A

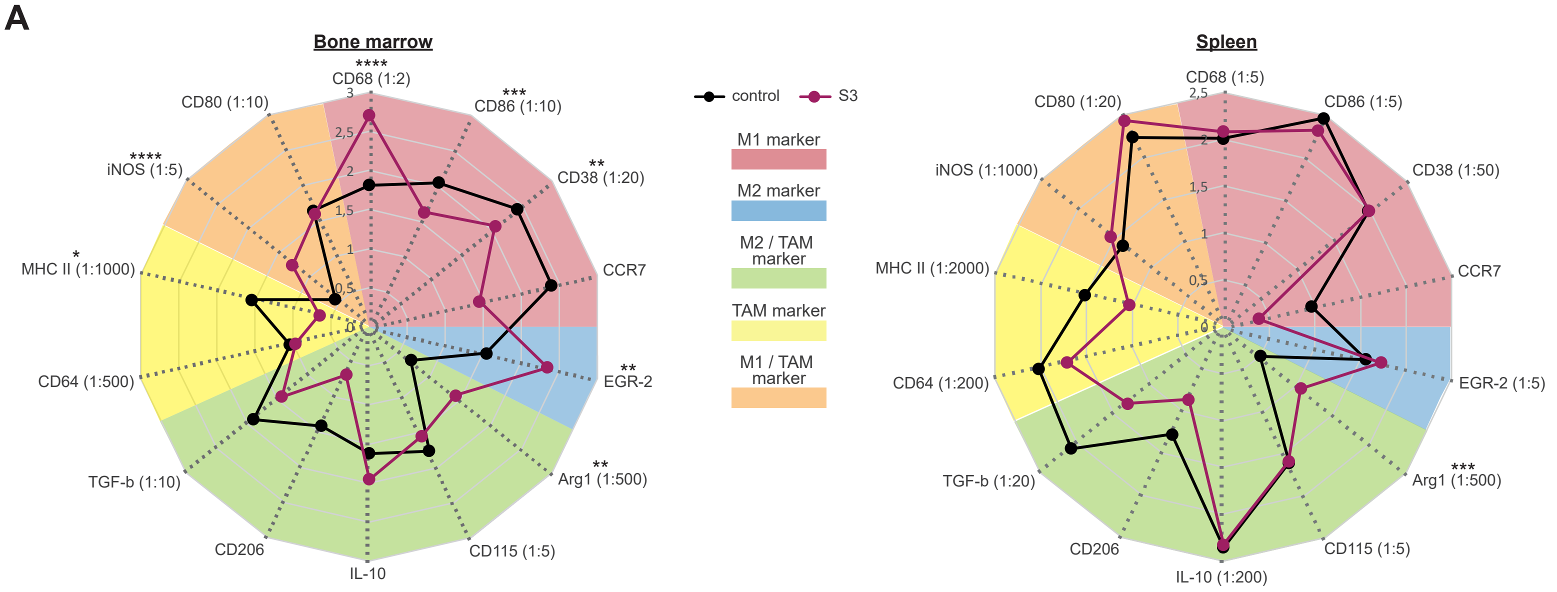


B

**Benjamini-Hochberg-analysis**

| | vehicle | alemtuzumab | S3 + alemtuzumab |
|---|---|---|---|
| vehicle | **--** | **1.4e-05** | **--** |
| S3 | **0.6159** | **5.5e-05** | **1.0e-07** |
| S3 + alemtuzumab | **3.4e-09** | **0.0059** | **--** |

C

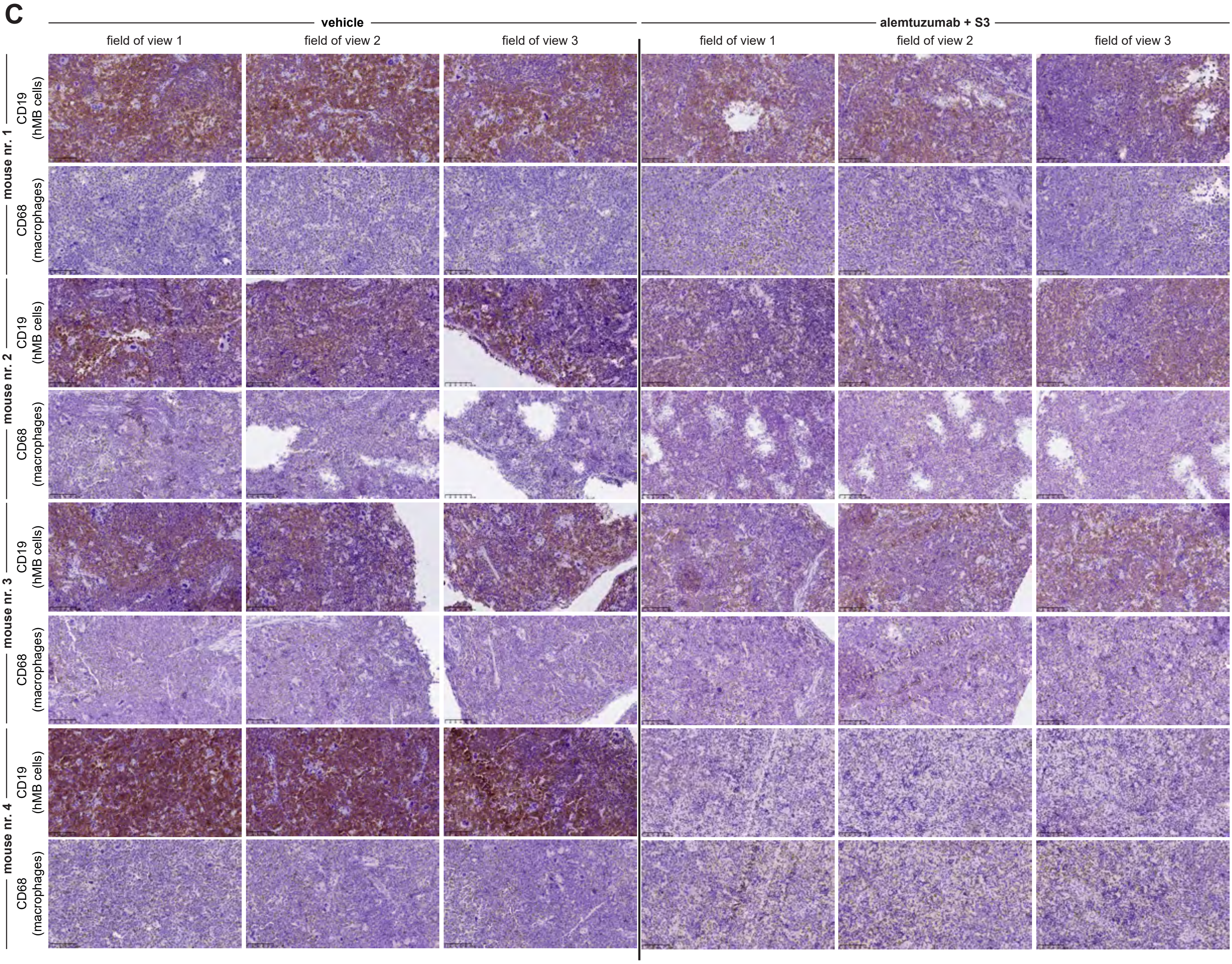

**Suppl. Fig.7 PPP inhibition increases myelopoiesis and macrophages´ activity in vivo and improves treatment response in an aggressive humanized lymphoma mouse model.**

**(A)** Expression of characteristic surface marker for different macrophage subtypes on macrophages in bone marrow and spleen. C57BL/6 mice treated with vehicle (control) or S3 i.p. for 7 days. **(B)** Significance testing by using Benjamini-Hochberg-analysis of survival curves of NSG mice transfected with hMB and treated after three days of engraftment with vehicle, alemtuzumab and/or PPP inhibitor S3 for 12 days. **(C)** Immunohistochemical staining of hMB cells ($CD19^{+}$) and macrophages ($CD68^{+}$) in spleen of NSG mice transfected with hMB and treated after three days of engraftment with vehicle or alemtuzumab + S3 for 12 days. **A** n=10, **B** n=21-25, **C** n=4. In **A** data are shown as mean of ten replicates. In **A** P values were calculated by using one-way ANOVA. $^{*}p < 0.05$; $^{**}p < 0.01$; $^{***}p < 0.001$; $^{****}p < 0.0001$.
